# The evolutionary origin of bilaterian smooth and striated myocytes

**DOI:** 10.1101/064881

**Authors:** Thibaut Brunet, Antje H. L. Fischer, Patrick R. H. Steinmetz, Antonella Lauri, Paola Bertucci, Detlev Arendt

## Abstract

The dichotomy between smooth and striated myocytes is fundamental for bilaterian musculature, but its evolutionary origin is unsolved. In particular, interrelationships of visceral smooth muscles remain unclear. Absent in fly and nematode, they have not yet been characterized molecularly outside vertebrates. Here, we characterize expression profile, ultrastructure, contractility and innervation of the musculature in the marine annelid Platynereis dumerilii and identify smooth muscles around the midgut, hindgut and heart that resemble their vertebrate counterparts in molecular fingerprint, contraction speed, and nervous control. Our data suggest that both visceral smooth and somatic striated myocytes were present in the protostome-deuterostome ancestor, and that smooth myocytes later co-opted the striated contractile module repeatedly – for example in vertebrate heart evolution. During these smooth-to-striated myocyte conversions the core regulatory complex of transcription factors conveying myocyte identity remained unchanged, reflecting a general principle in cell type evolution.

## Introduction

Musculature is composed of myocytes that are specialized on active contraction (Schmidt-Rhaesa 2007). Their contractile apparatus centers on actomyosin, a contractile module that dates back to stem eukaryotes (Brunet & Arendt 2016) and incorporated accessory proteins of pre-metazoan origin (Steinmetz et al. 2012). Two fundamentally distinct types of myocytes are distinguished based on ultrastructural appearance. In striated myocytes, actomyosin myofibrils are organized in aligned repeated units (sarcomeres) separated by transverse ‘Z discs’, while in smooth myocytes adjacent myofibrils show no clear alignment and are separated by scattered “dense bodies” (Figure 1A). In vertebrates, striated myocytes are found in voluntary skeletal muscles, but also at the anterior and posterior extremities of the digestive tract (anterior esophagus muscles and external anal sphincter), and in the muscular layer of the heart; smooth myocytes are found in involuntary visceral musculature that ensures slow, long-range deformation of internal organs. This includes the posterior esophagus and the rest of the gut, but also blood vessels, and most of the urogenital system. In stark contrast, in the fruit fly *Drosophila* virtually all muscles are striated, including gut visceral muscles (Anderson & Ellis 1967; Goldstein & Burdette 1971; Paniagua et al. 1996); the only exception are poorly-characterized multinucleated smooth muscles around the testes (Susic-Jung et al. 2012). Also, in the nematode *Caenorhabditis*, somatic muscles are striated, while the short intestine and rectum visceral myocytes are only one sarcomere-long and thus hard to classify (White 1988; Corsi et al. 2000).

**Figure 1.**
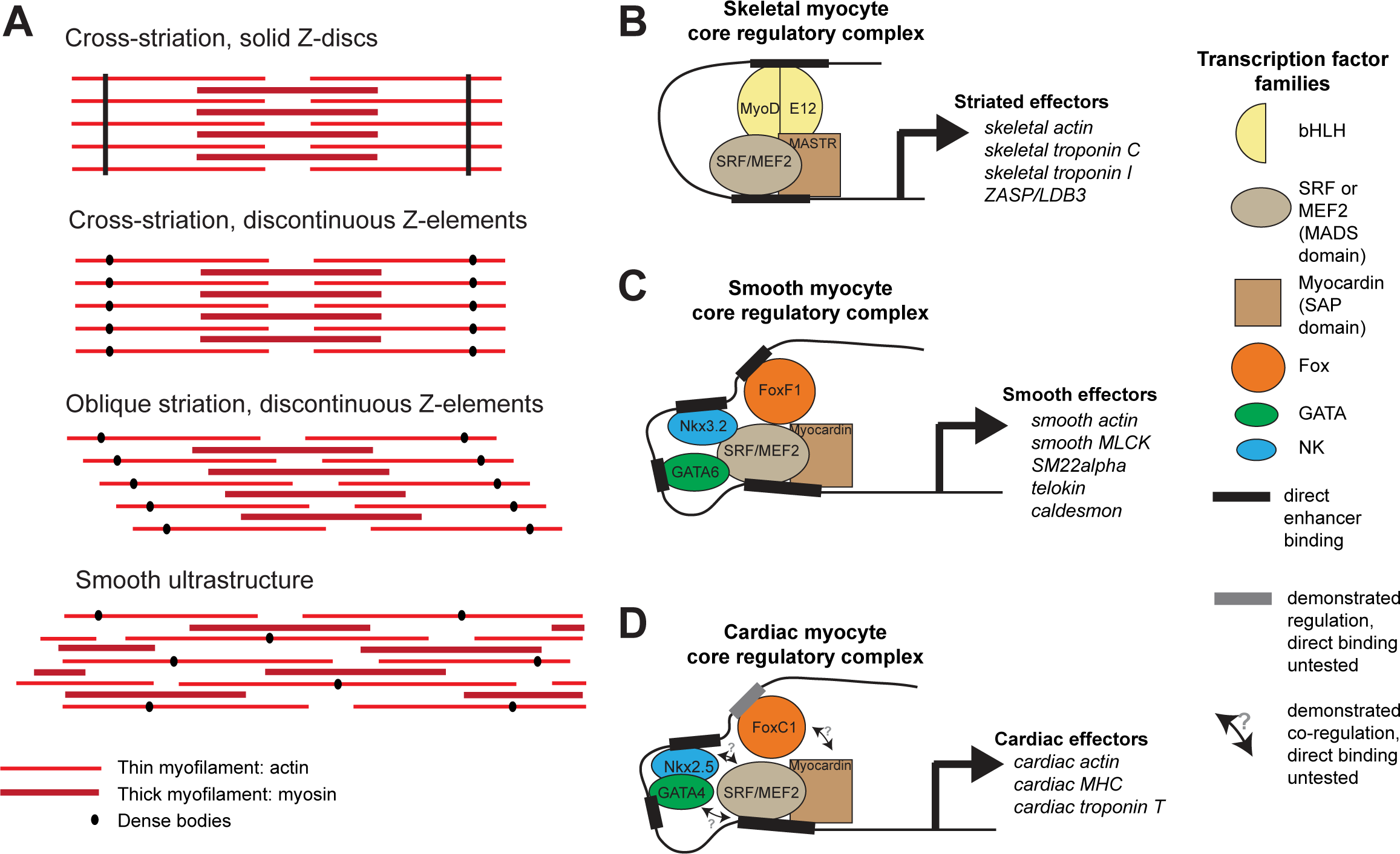
Ultrastructure and core regulatory complexes of myocyte types. (A) Schematic smooth and striated ultrastructures. Electron-dense granules called “dense bodies” separate adjacent myofibrils. Dense bodies are scattered in smooth muscles, but aligned in striated muscles to form Z lines. (B-D) Core regulatory complexes (CRC) of transcription factors for the differentiation of different types of myocytes in vertebrates. Complexes composition from (Creemers et al. 2006; Meadows et al. 2008; Molkentin et al. 1995) for skeletal myocytes, (Hoggatt et al. 2013; Nishida et al. 2002; Phiel et al. 2001) for smooth myocytes, and (Durocher et al. 1997; Lee et al. 1998) for cardiomyocytes. Target genes from (Blais et al. 2005) for skeletal myocytes, (Nishida et al. 2002) from smooth myocytes, and (Schlesinger et al. 2011) for cardiomyocytes.

The evolutionary origin of smooth versus striated myocytes in bilaterians accordingly remains unsolved. Ultrastructural studies have consistently documented the presence of striated somatic myocytes in virtually every bilaterian group (Schmidt-Rhaesa 2007) and in line with this, the comparison of Z-disc proteins supports homology of striated myocytes across bilaterians (Steinmetz et al. 2012). The origin of smooth myocyte types however is less clear. Given the absence of smooth muscles from fly and nematode, it has been proposed that visceral smooth myocytes represent a vertebrate novelty, which evolved independently from non-muscle cells in the vertebrate stem line (Goodson & Spudich 1993; OOta & Saitou 1999). However, smooth muscles are present in many other bilaterian groups, suggesting instead their possible presence in urbilaterians and secondary loss in arthropods and nematodes. Complicating the matter further, intermediate ultrastructures between smooth and striated myocytes have been reported, suggesting interconversions (reviewed in (Schmidt-Rhaesa 2007))

Besides ultrastructure, the comparative molecular characterization of cell types can be used to build cell type trees (Arendt 2003; Arendt 2008; Wagner 2014; Musser & Wagner 2015). Cell type identity is established via the activity of transcription factors acting as terminal selectors (Hobert 2016) and forming “core regulatory complexes” (CRCs; (Arendt et al. 2016; Wagner 2014)), which directly activate downstream effector genes. This is exemplified for vertebrate myocytes in Figure 1B. In all vertebrate myocytes, transcription factors of the Myocardin family (MASTR in skeletal muscles, Myocardin in smooth and cardiac muscles) directly activate effector genes encoding contractility proteins (Fig. 1B) (Creemers et al. 2006; Meadows et al. 2008; Wang & Olson 2004; Wang et al. 2003)). They heterodimerize with MADS-domain factors of the Myocyte Enhancer Factor-2 (Mef2) (Black & Olson 1998; Blais et al. 2005; Molkentin et al. 1995; Wales et al. 2014) and Serum Response Factor (SRF) families (Carson et al. 1996; Nishida et al. 2002). Other myogenic transcription factors are specific for different types of striated and smooth myocytes. Myogenic Regulatory Factors (MRF) family members, including MyoD and its paralogs Myf5, Myogenin, and Mrf4/Myf6 (Shi & Garry 2006), directly control contractility effector genes in skeletal (and esophageal) striated myocytes, cooperatively with Mef2 (Blais et al. 2005; Molkentin et al. 1995) – but are absent from smooth and cardiac muscles. In smooth and cardiac myocytes, this function is ensured by NK transcription factors (Nkx3.2/Bapx and Nkx2.5/Tinman, respectively), GATA4/5/6, and Fox transcription factors (FoxF1 and FoxC1, respectively), which bind to SRF and Mef2 to form CRCs directly activating contractility effector genes (Durocher et al. 1997; Hoggatt et al. 2013; Lee et al. 1998; Morin et al. 2000; Nishida et al. 2002; Phiel et al. 2001) (Figure 1B).

Regarding effector proteins (Figure 1B), (Kierszenbaum & Tres 2015), all myocytes express distinct isoforms of the myosin heavy chain: the striated myosin heavy chain *ST-MHC* (which duplicated into cardiac, fast skeletal, and slow skeletal isoforms in vertebrates) and the smooth/non-muscle myosin heavy chain *SM-MHC* (which duplicated in vertebrates into smooth *myh10, myh11* and *myh14*, and non-muscle *myh9*) (Steinmetz et al. 2012). The different contraction speeds of smooth and striated muscles are due to the distinct kinetic properties of these molecular motors (Bárány 1967). In both myocyte types, contraction occurs in response to calcium, but the responsive proteins differ (Alberts et al. 2014): the Troponin complex (composed of Troponin C, Troponin T and Troponin I) for striated muscles, Calponin and Caldesmon for smooth muscles. In both myocyte types, calcium also activates the Calmodulin/Myosin Light Chain Kinase pathway (Kamm & Stull 1985; Sweeney et al. 1993). Striation itself is implemented by specific effectors, including the long elastic protein Titin (Labeit & Kolmerer 1995) (which spans the entire sarcomere and confers it elasticity and resistance) and ZASP/LBD3 (Z-band Alternatively Spliced PDZ Motif/LIM-Binding Domain 3), which binds actin and stabilizes sarcomeres during contraction (Au et al. 2004; Zhou et al. 2001). The molecular study of *Drosophila* and *Caenorhabditis* striated myocytes revealed important commonalities with their vertebrate counterparts, including the Troponin complex (Beall & Fyrberg 1991; Fyrberg et al. 1994; Fyrberg et al. 1990; Marín et al. 2004; Myers et al. 1996), and a conserved role for Titin (Zhang et al. 2000) and ZASP/LBD3 (Katzemich et al. 2011; McKeown et al. 2006) in the striated architecture.

Finally, smooth and striated myocytes also differ physiologically. All known striated myocyte types (apart from the myocardium) strictly depend on nervous stimulations for contraction, exerted by innervating motor neurons. In contrast, gut smooth myocytes are able to generate and propagate automatic (or “myogenic”) contraction waves responsible for digestive peristalsis in the absence of nervous inputs (Faussone - Pellegrini & Thuneberg 1999; Sanders et al. 2006). These autonomous contraction waves are modulated by the autonomic nervous system (Silverthorn 2015). Regarding overall contraction speed, striated myocytes have been measured to contract 10 to 100 times faster than their smooth counterparts (Bárány 1967).

To elucidate the evolutionary origin and diversification of bilaterian smooth and striated myocytes, we provide an in-depth ultrastructural, molecular and functional characterization of the myocyte complement in the marine annelid *Platynereis dumerilii*, which belongs to the Lophotrochozoa. Strikingly, as of now, no invertebrate smooth visceral muscle has been investigated on a molecular level (Hooper & Thuma 2005; Hooper et al. 2008). *Platynereis* has retained more ancestral features than flies or nematodes and is thus especially suited for long-range comparisons (Raible et al. 2005; Denes et al. 2007). Also, other annelids such as earthworms have been reported to possess both striated somatic and midgut smooth visceral myocytes based on electron microscopy (Anderson & Ellis 1967). Our study reveals the parallel presence of smooth myocytes in the musculature of midgut, hindgut, and pulsatile dorsal vessel and of striated myocytes in the somatic musculature and the foregut. *Platynereis* smooth and striated myocytes closely parallel their vertebrate counterparts in ultrastructure, molecular profile, contraction speed, and reliance on nervous inputs, thus supporting the ancient existence of a smooth-striated duality in protostome/deuterostome ancestors.

## Results

### Platynereis midgut and hindgut muscles are smooth, while foregut and somatic muscles are striated

Differentiation of the *Platynereis* somatic musculature has been documented in much detail (Fischer et al. 2010) and, in five days old young worms, consists of ventral and dorsal longitudinal muscles, oblique and parapodial muscles, head muscles and the axochord (Lauri et al. 2014). At this stage, the first *Platynereis* visceral myocytes become detectable around the developing tripartite gut, which is subdivided into foregut, midgut and hindgut (based on the conserved regional expression of *foxA, brachyury* and *hnf4* gut specification factors (Martín-Durán & Hejnol 2015); Supplementary Figure 1). At 7 days post fertilization (dpf), visceral myocytes form circular myofibres around the foregut, and scattered longitudinal and circular fibres around midgut and hindgut (Figure 2A), which expand by continuous addition of circular and longitudinal fibres to completely cover the dorsal midgut at 11dpf (Figure 2A) and finally form a continuous muscular orthogon around the entire midgut and hindgut in the 1.5 months-old juvenile (Figure 2A).

**Figure 2.**
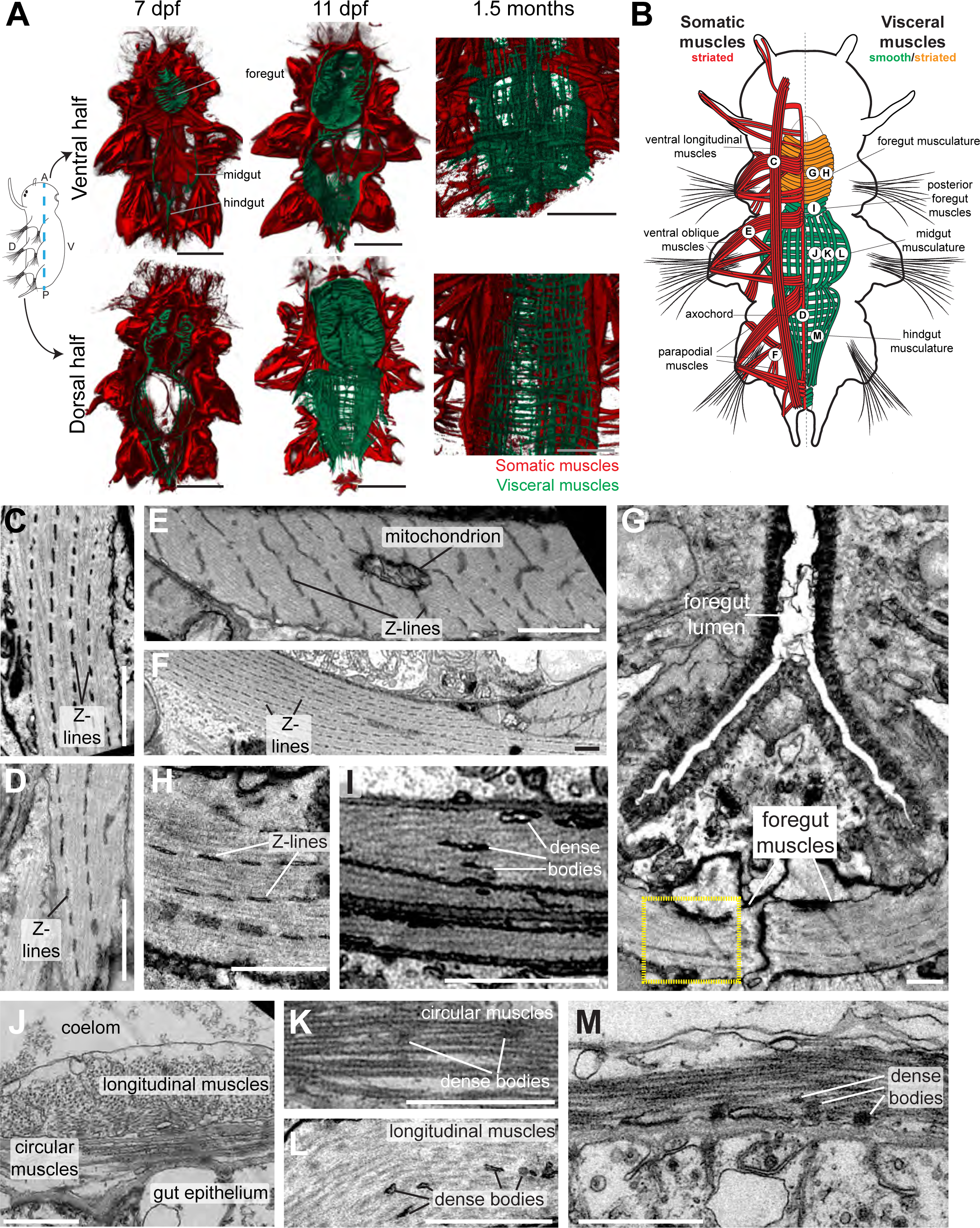
Development and ultrastrcture of visceral and somatic musculature in *Platynereis* larvae and juveniles. (A) Development of visceral musculature. All panels are 3D renderings of rhodamine-phalloidin staining imaged by confocal microscopy. Visceral muscles have been manually colored green and somatic muscle red. Scale bar: 50 µm. (B) Schematic of the musculature of a late nectochaete (6 dpf) larva. Body outline modified from (Fischer et al. 2010). Ventral view, anterior is up. (C-M) Electron micrographs of the main muscle groups depicted in B. Each muscle group is shown sectioned parallel to its long axis, so in the plane of its myofilaments. Scale bar: 2 µm. (H) is a close-up of the region in the yellow box in G. (J) shows the dorsal midgut in cross-section, dorsal side up.

We then proceeded to characterize the ultrastructure of *Platynereis* visceral and somatic musculature by transmission electron microscopy (Figure 2C-M). All somatic muscles and anterior foregut muscles display prominent oblique striation with discontinuous Z-elements (Figure 2C-H; compare Figure 1A), as typical for protostomes (Burr & Gans 1998; Mill & Knapp 1970; Rosenbluth 1972). To the contrary, visceral muscles of the posterior foregut, midgut and hindgut are smooth with scattered dense bodies (Figure 2I-M). The visceral muscular orthogon is partitioned into an external longitudinal layer and an internal circular layer (Figure 2J), as in vertebrates (Marieb & Hoehn 2015) and arthropods (Lee et al. 2006). Thus, according to ultrastructural appearance, *Platynereis* has both somatic (and anterior foregut) striated muscles and visceral smooth muscles.

### The molecular profile of smooth and striated myocytes

We then set out to molecularly characterize annelid smooth and striated myocytes via a candidate gene approach. As a starting point, we investigated, in the *Platynereis* genome, the presence of regulatory and effector genes specific for smooth and/or striated myocytes in the vertebrates. We found striated muscle-specific and smooth muscle/non-muscle isoforms of both *myosin heavy chain* (consistently with published phylogenies (Steinmetz et al. 2012)) and *myosin regulatory light chain*. We also identified homologs of genes encoding calcium transducers (*calponin* for smooth muscles; *troponin I* and *troponin T* for striated muscles), striation structural proteins (*zasp/lbd3* and titin), and terminal selectors for the smooth (*nk3, foxF*, and *gata456*) and striated phenotypes *(myoD)*.

We investigated expression of these markers by whole-mount in situ hybridization (WMISH). Striated effectors are expressed in both somatic and foregut musculature (Figure 3A, C; Figure 3—supplement 1). Expression of all striated effectors was observed in every somatic myocyte group by confocal imaging with cellular resolution (Figure 3— supplement 2). Interestingly, *myoD* is exclusively expressed in longitudinal striated muscles, but not in other muscle groups (Figure 3—supplement 2).

**Figure 3.**
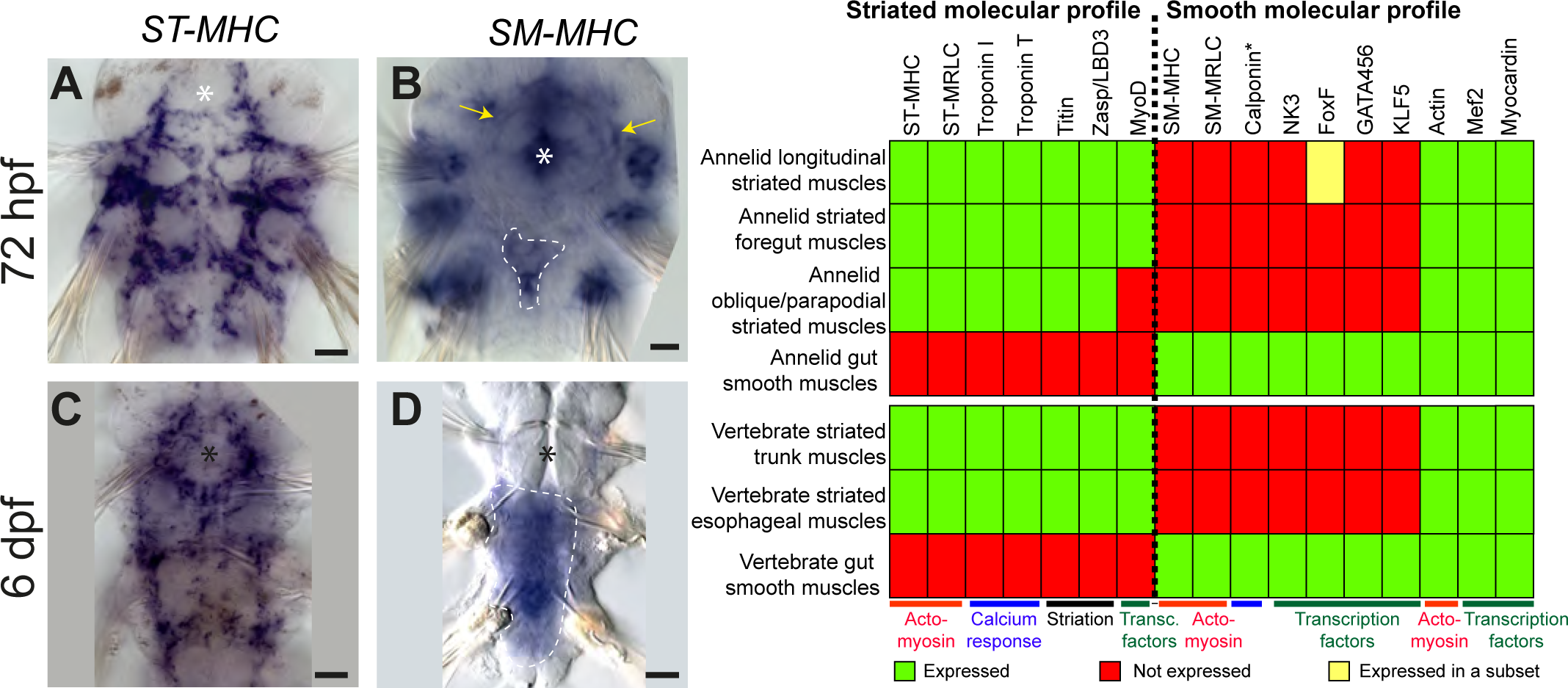
Expression of smooth and striated muscle markers in *Platynereis* larvae. Animals have been stained by WMISH and observed in bright field Nomarski microscopy. Ventral views, anterior side up. Scale bar: 25 µm. (A-D) Expression patterns of the striated marker *ST-MHC* and the smooth marker *SM-MHC*. These expression patterns are representative of the entire striated and smooth effector module (see Supplementary Figures 2 and 4). (E) Table summarizing the expression patterns of smooth and striated markers in *Platynereis* and vertebrate muscles. (*) indicates that *Platynereis* and vertebrate Calponin are mutually most resembling by domain structure, but not one-to-one orthologs, as independent duplications in both lineages have given rise to more broadly expressed paralogs with a different domain structure (Figure 7—supplement 3).

The expression of smooth markers is first detectable at 3 dpf in a small triangle-shaped group of mesodermal cells posteriorly abutting the macromeres (which will form the future gut) (Figure 3B, Figure 3—supplement 3A-D, H). Double WMISH shows that these cells coexpress smooth markers and the muscle marker *actin* (Figure 3— supplement 3I-K), supporting the idea that they are forming gut myocytes. At this stage, smooth markers are also expressed in the foregut mesoderm (Figure 3B, Figure 3— supplement 3A-D,H, yellow arrows). At 6 dpf, expression of all smooth markers (apart from *nk3*) is maintained in the midgut and hindgut differentiating myocytes (Figure 3D, Figure 3—supplement 3E-G, Figure 3—supplement 4A-E) but smooth effectors disappear from the foregut, which turns on striated markers instead (Figure 3— supplement 1R-W) – reminiscent of the replacement of smooth fibres by striated fibres during development of the vertebrate anterior esophageal muscles (Gopalakrishnan et al. 2015). Finally, in 2 months-old juvenile worms, smooth markers are also detected in the dorsal pulsatile vessel (Figure 3—supplement 3L-Q) – considered equivalent to the vertebrate heart (Saudemont et al. 2008) but, importantly, of smooth ultrastructure in polychaetes (Spies 1973; Jensen 1974). None of the striated markers is expressed around the midgut or the hindgut (Figure 3—supplement 4F-K), or in the dorsal vessel (Figure 3—supplement 3P). Taken together, these results strongly support conservation of the molecular fingerprint of both smooth and striated myocytes between annelids and vertebrates.

We finally investigated general muscle markers that are shared between smooth and striated muscles. These include *actin, mef2* and *myocardin* – which duplicated into muscle type-specific paralogs in vertebrates, but are still present as single-copy genes in *Platynereis*. We found them to be expressed in the forming musculature throughout larval development (Figure 3—supplement 5A-F), and confocal imaging at 6 dpf confirmed expression of all 3 markers in both visceral (Figure 3—supplement 5G-L) and somatic muscles (Figure 3—supplement 5M).

### Smooth and striated muscles differ in contraction speed

We then characterized the contraction speed of the two myocyte types in *Platynereis* by measuring myofibre length before and after contraction. Live confocal imaging of contractions in *Platynereis* larvae with fluorescently labeled musculature (Movie 1, Movie 2) gave a striated contraction rate of 0.55±0.27 s^−1^ (Figure 4A-E) and a smooth myocyte contraction rate of 0.07±0.05 s^−1^ (Figure 4G). As in vertebrates, annelid striated myocytes thus contract nearly one order of magnitude faster than smooth myocytes (Figure 4F).

**Figure 4.**
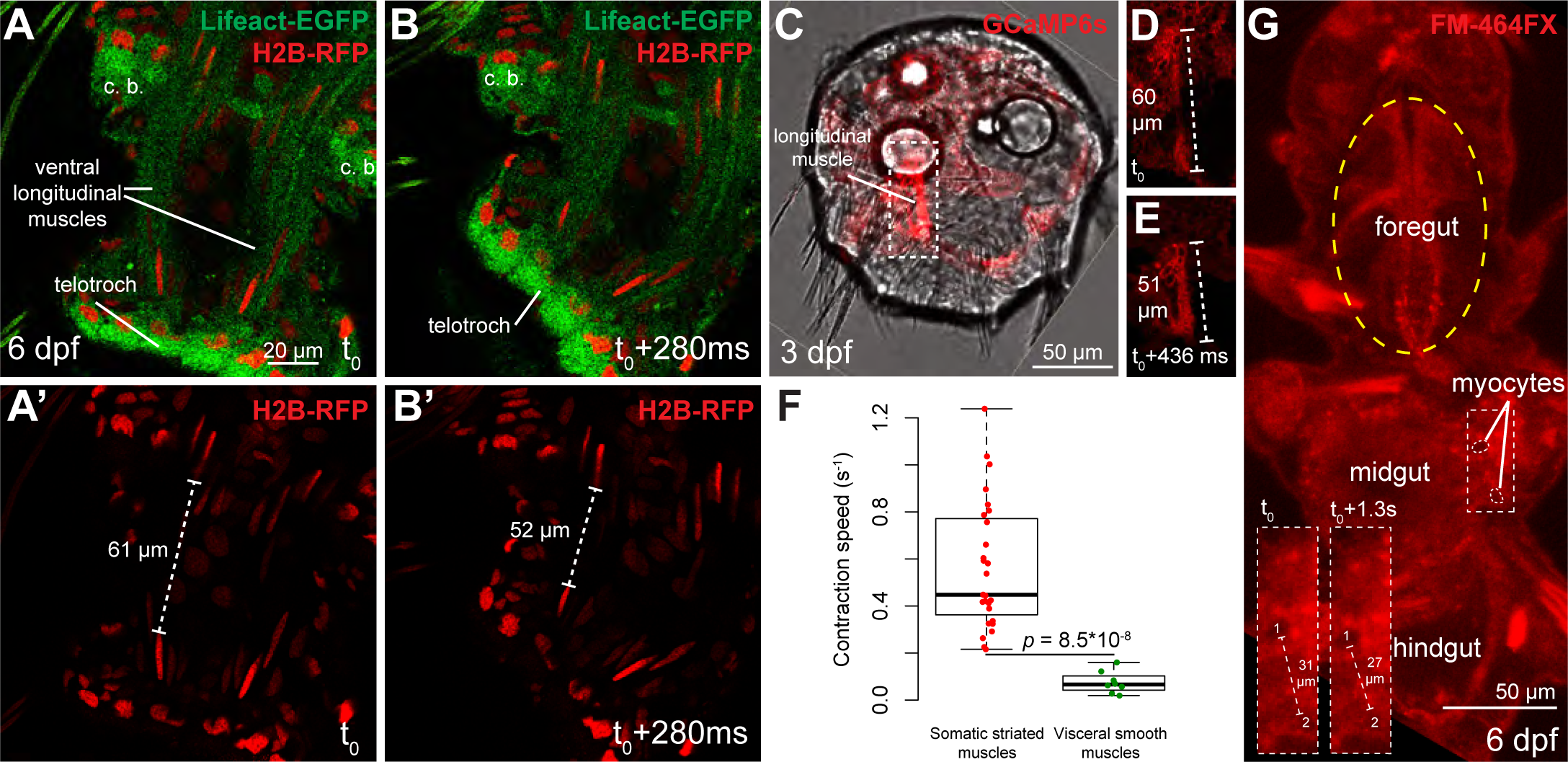
Contraction speed quantifications of smooth and striated muscles. (AB) Snapshots of a time lapse live confocal imaging of a late nectochaete larva expressing fluorescent markers. Ventral view of the 2 posterior-most segments, anterior is up. (C) Snapshots of a time lapse live confocal imaging of a 3 dpf larva expressing GCaMP6s. Dorsal view, anterior is up. (D-E) Two consecutive snapshots on the left dorsal longitudinal muscle of the larva shown in C, showing muscle contraction. (F) Quantification of smooth and striated muscle contraction speeds (see Experimental procedures and Figure 4-Source data 1), p-value by Mann-Whitney’s U test. Each point represents a biological replicate (see Material and Methods). (G) Snapshot of a time lapse live confocal imaging of a late nectochaete larva. Ventral view, anterior is up. Optical longitudinal section at the midgut level.

### Striated, but not smooth, muscle contraction depends on nervous inputs

Finally, we investigated the nervous control of contraction of both types of muscle cells. In vertebrates, somatic muscle contraction is strictly dependent on neuronal inputs. By contrast, gut peristalsis is automatic (or myogenic – i.e., does not require nervous inputs) in vertebrates, cockroaches (Nagai & Brown 1969), squids (Wood 1969), snails (Roach 1968), holothurians, and sea urchins (Prosser et al. 1965). The only exceptions appear to be bivalves and malacostracans (crabs, lobster and crayfish), in which gut motility is neurogenic (Prosser et al. 1965). Regardless of the existence of an automatic component, the gut is usually innervated by nervous fibres modulating peristalsis movements (Wood 1969; Wu 1939).

Gut peristalsis takes place in *Platynereis* larvae and juveniles from 6 dpf onwards (Movie 3), and we set out to test whether nervous inputs were necessary for it to take place. We treated 2 months-old juveniles with 180 μM Brefeldin A, an inhibitor of vesicular traffic which prevents polarized secretion (Misumi et al. 1986) and neurotransmission (Malo et al. 2000). Treatment stopped locomotion in all treated individuals, confirming that neurotransmitter release by motor neurons is required for somatic muscles contraction, while DMSO-treated controls were unaffected. On the other hand, vigorous gut peristalsis movements were maintained in Brefeldin A-treated animals (Movie 4). Quantification of the propagation speed of the peristalsis wave (Figure 5A-D; see Material and Methods) indicated that contractions propagated significantly faster in Brefeldin A-treated individuals than in controls. The frequency of wave initiation and their recurrence (the number of repeated contraction waves occurring in one uninterrupted sequence) did not differ significantly in Brefeldin A-treated animals (Figure 5E,F). These results indicate that, as in vertebrates, visceral smooth muscle contraction and gut peristalsis do not require nervous (or secretory) inputs in *Platynereis*.

**Figure 5.**
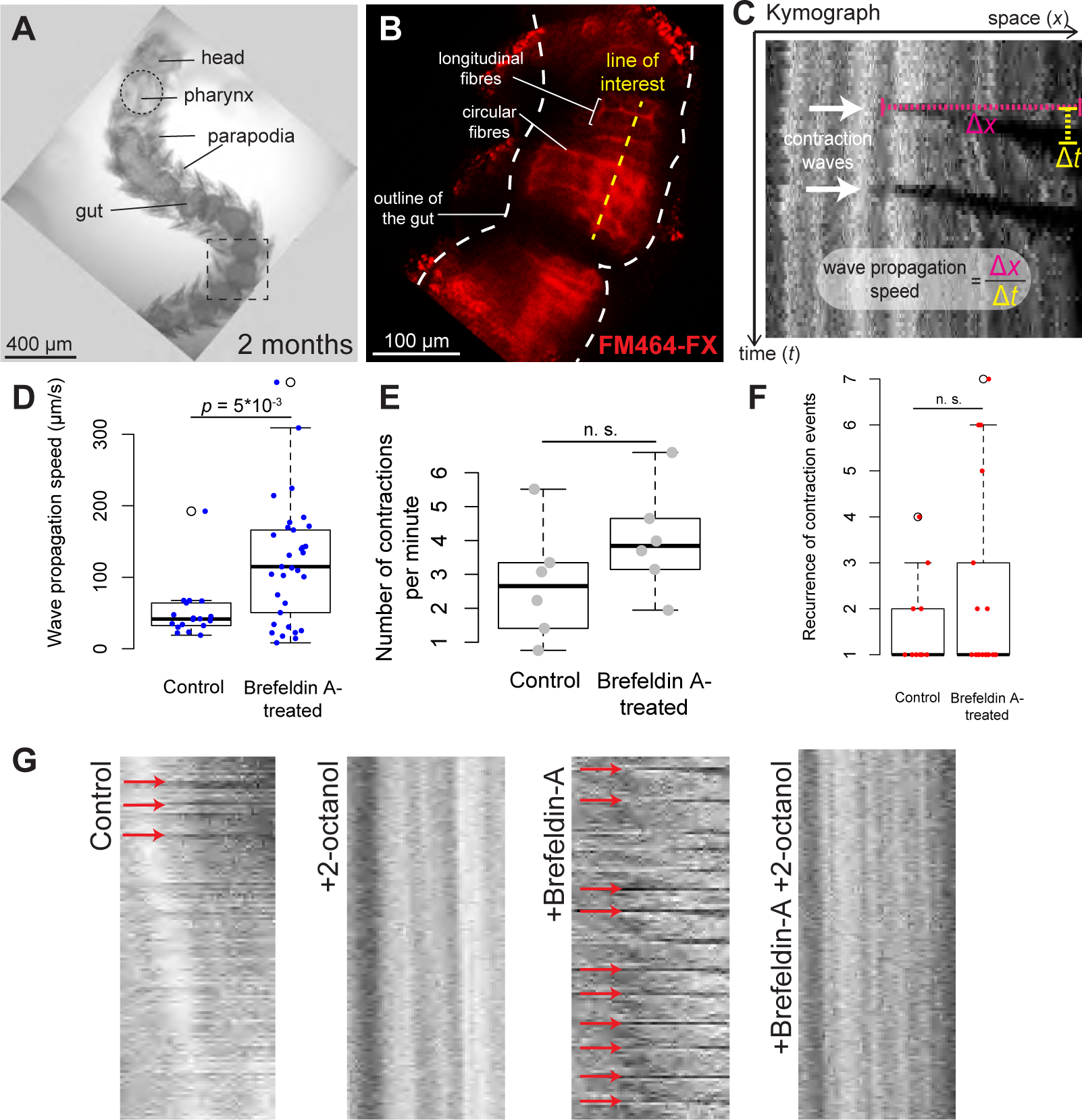
*Platynereis* gut peristalsis is independent of nervous inputs and dependent on gap junctions. (A) 2 months-old juvenile mounted in 3% low-melting point (LMP) agarose for live imaging. (B) Snapshots of a confocal live time lapse imaging of the animal shown in A. Gut is observed by detecting fluorescence of the vital membrane dye FM-464FX. (C) Kymograph of gut peristalsis along the line of interest in (B). Contraction waves appear as dark stripes. A series of consecutive contraction waves is called a *contraction event*: here, two contraction waves are visible, which make up one contraction event with a recurrence of 2. (D) Quantification of the propagation speed of peristaltic contraction waves in mock (DMSO)-treated individuals and Brefeldin A-treated individuals (inhibiting neurotransmission). Speed is calculated from kymographs (see Material and Methods and Figure 5-source data 1), *p*-value by Mann-Whitney’s U test. Each point represents a contraction wave. 5 biological replicates for each category (see Material and Methods). (E, F) Same as in E, but showing respectively the frequency of initiation and the recurrence of contraction events. Each point represents a biological replicate (see Material and Methods).(G) Representative kymographs of controls, animals treated with Brefeldin A (inhibiting neurotransmission), animals treated with 2-octanol (inhibiting gap junctions), and animals treated with both (N= 10 for each condition). 2-octanol entirely abolishes peristaltic waves with or without Brefeldin A.

### An enteric nervous system is present in Platynereis

In vertebrates, peristaltic contraction waves are initiated by self-excitable myocytes (Interstitial Cajal Cells) and propagate across other smooth muscles by gap junctions ensuring direct electrical coupling (Faussone - Pellegrini & Thuneberg 1999; Sanders et al. 2006). We tested the role of gap junctions in *Platynereis* gut peristalsis by treating animals with 2.5 mM 2-octanol, which inhibits gap junction function in both insects (Bohrmann & Haas-Assenbaum 1993; Gho 1994) and vertebrates (Finkbeiner 1992). 2-octanol abolishes gut peristalsis, both in the absence and in the presence of Brefeldin A (Figure 5G), indicating that propagation of the peristalsis wave relies on direct coupling between smooth myocytes via gap junctions.

The acceleration of peristalsis upon Brefeldin A treatment suggests that secretory (and possibly neural) inputs modulate gut contractions (with a net inhibitory effect). We thus investigated the innervation of the *Platynereis* gut. Immunostainings of juvenile worms for acetylated tubulin revealed a dense, near-orthogonal nerve net around the entire gut (Figure 6A), which is tightly apposed to the visceral muscle layer (Figure 6C) and includes serotonergic neurites (Figure 6B,C) and cell bodies (Figure 6D). Interestingly, some enteric serotonergic cell bodies are devoid of neurites, thus resembling the vertebrate (non-neuronal) enterochromaffine cells – endocrine serotonergic cells residing around the gut and activating gut peristalsis by direct serotonin secretion upon mechanical stretch (Bulbring & Crema 1959).

**Figure 6.**
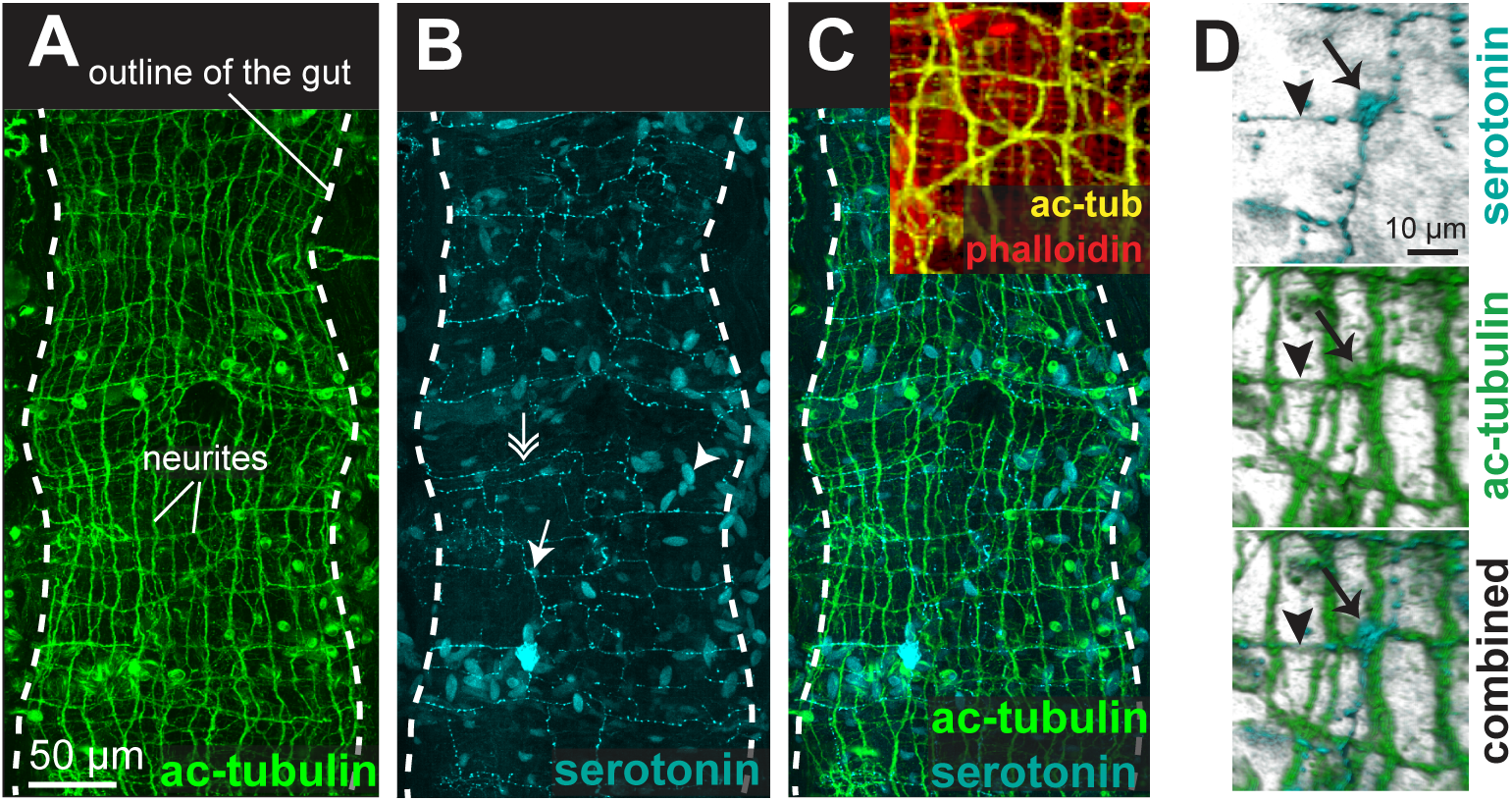
The enteric nerve net of *Platynereis*. (A) Immunostaining for acetylated tubulin, visualizing neurites of the enteric nerve plexus. Z-projection of a confocal stack at the level of the midgut. Anterior side up. (B) Same individual as in A, immunostaining for serotonin (5-HT). Note serotonergic neurites (double arrow), serotonergic neuronal cell bodies (arrow, see D), and serotonergic cell bodies without neurites (arrowhead). (C) Same individual as in A showing both acetylated tubulin and 5-HT immunostainings. Snapshot in the top right corner: same individual, showing both neurites (acetylated tubulin, yellow) and visceral myofibres (rhodamine-phalloidin, red). The acetylated tubulin appears yellow due to fluorescence leaking in the rhodamine channel. (D) 3D rendering of the serotonergic neuron shown by arrow in B.

## Discussion

### Smooth and striated myocyte coexisted in bilaterian ancestors

Our study represents the first molecular characterization of protostome visceral smooth musculature (Hooper & Thuma 2005; Hooper et al. 2008). The conservation of molecular signatures for both smooth and striated myocytes indicates that a dual musculature already existed in bilaterian ancestors: a fast striated somatic musculature (possibly also present around the foregut – as in *Platynereis*, vertebrates (Gopalakrishnan et al. 2015) and sea urchins (Andrikou et al. 2013; Burke 1981)), under strict nervous control; and a slow smooth visceral musculature around the midgut and hindgut, able to undergo automatic peristalsis thanks to self-excitable myocytes directly coupled by gap junctions, and undergoing modulation by nervous and paracrine inputs. In striated myocytes, a core regulatory complex (CRC) involving Mef2 and Myocardin directly activated striated contractile effector genes such as *ST-MHC*, *ST-MRLC* and the *Troponin* genes (Figure 7—supplement 1). Notably, myoD might have been part of the CRC in only part of the striated myocytes, as it is only detected in longitudinal muscles in *Platynereis*. The absence of *myoD* expression in other annelid muscle groups is in line with the “chordate bottleneck” concept (Thor & Thomas 2002), according to which specialization for undulatory swimming during early chordate evolution would have fostered exclusive reliance on trunk longitudinal muscles, and loss of other muscle types. In smooth myocytes, a CRC composed of NK3, FoxF and GATA4/5/6 together with Mef2 and Myocardin activated the smooth contractile effectors *SM-MHC*, *SM-MRLC* and *calponin* (Figure 7—supplement 1). In spite of their absence in flies and nematodes, gut myocytes of smooth ultrastructure are widespread in other bilaterians, and an ancestral state reconstruction retrieves them as present in the last common protostome/deuterostome ancestor with high confidence (Figure 7—supplement 2), supporting our homology hypothesis. Our results are consistent with previous reports of Calponin immunoreactivity in intestinal muscles of earthworms (Royuela et al. 1997) and snails (which also lack immunoreactivity for Troponin T) (Royuela et al. 2000).

### Origin of smooth and striated myocytes by cell type individuation

How did smooth and striated myocytes diverge in evolution? Figure 7 presents a comprehensive cell type tree for the evolution of myocytes, with a focus on Bilateria. This tree illustrates the divergence of the two muscle cell types by progressive partitioning of genetic information in evolution – a process called *individuation* (Wagner 2014; Arendt et al. 2016). The individuation of fast and slow contractile cells involved two complementary processes: (1) changes in CRC (black circles, Figure 7) and (2) emergence of novel genes encoding new cellular modules, or *apomeres* (Arendt et al. 2016) (grey squares, Figure 7).

**Figure 7.**
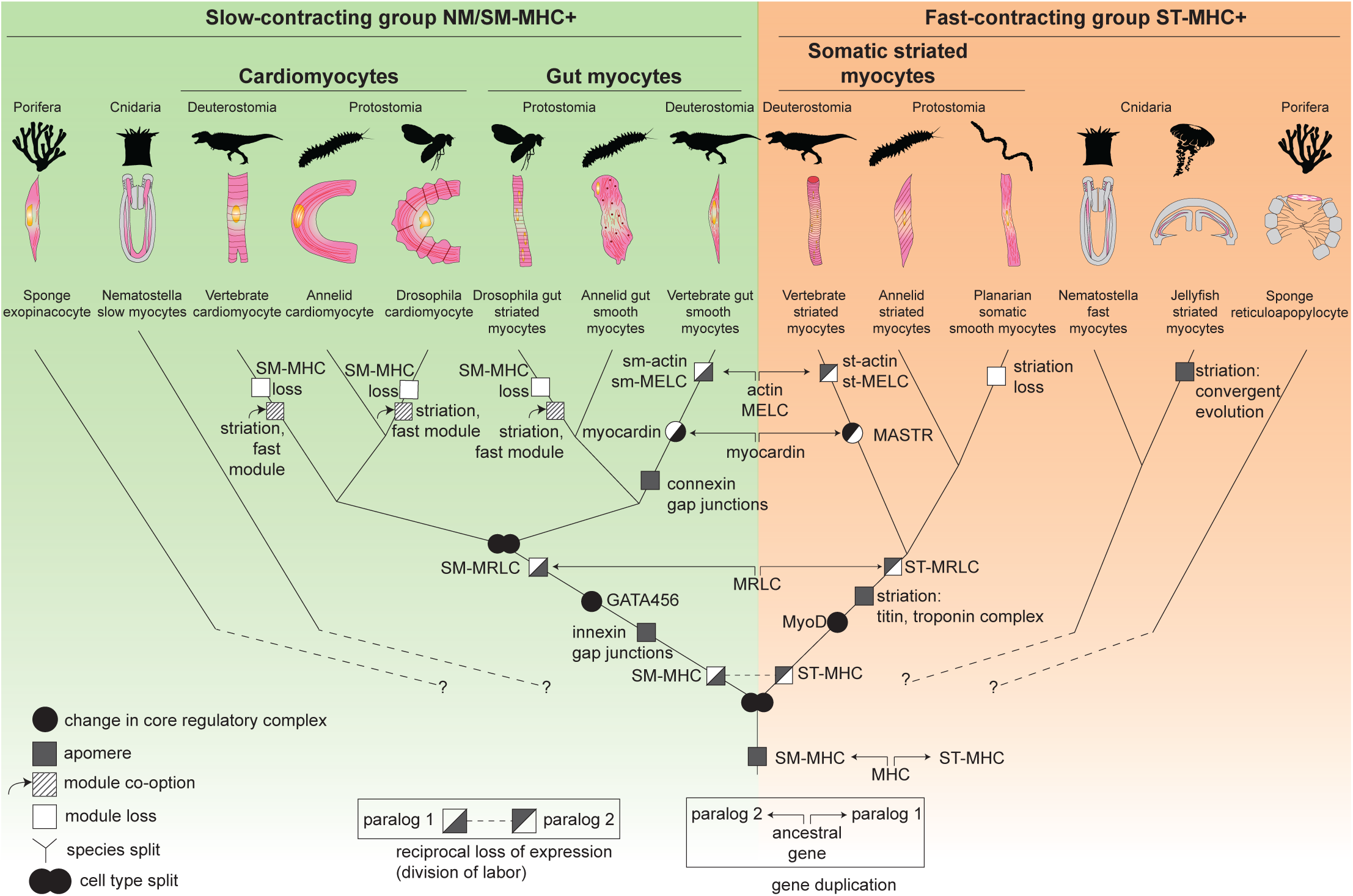
The evolutionary tree of animal contractile cell types. Bilaterian smooth and striated muscles split before the last common protostome/deuterostome ancestor. Bilaterian myocytes are split into two monophyletic cell type clades: an ancestrally *SM­ MHC+* slow-contracting clade (green) and an ancestrally *ST-MHC+* fast-contracting clade (orange). Hypothetical relationships of the bilaterian myocytes to the *SM-MHC+* and *ST-MHC*+ contractile cells of non-bilaterians are indicated by dotted lines (Steinmetz et al. 2012). Apomere: derived set of effector genes common to a monophyletic group of cell types (Arendt et al. 2016). Note that ultrastructure only partially reflects evolutionary relationships, as striation can evolve convergently (as in medusozoans), be co-opted (as in insect gut myocytes or in vertebrate and insect cardiomyocytes), be blurred, or be lost (as in planarians). Conversion of smooth to striated myocytes took place by co-option of striation proteins (Titin, Zasp/LDB3) and of the fast contractile module (ST-MHC, ST­ MRLC, Troponin complex) in insect cardiomyocytes and gut myocytes, as well as in vertebrate cardiomyocytes.

Around a common core formed by the Myocardin:Mef2 complex (both representing transcription factors of pre-metazoan ancestry (Steinmetz et al. 2012)), smooth and striated CRCs incorporated different transcription factors implementing the expression of distinct effectors (Figure 1B; Figure 7—supplement 1) – notably the bilaterian-specific bHLH factor MyoD (Steinmetz et al. 2012) and GATA4/5/6, which arose by bilaterian-specific duplication of a single ancient pan-endomesodermal GATA transcription factor (Martindale et al. 2004; Leininger et al. 2014).

Regarding the evolution of myocyte-specific apomeres, one prominent mechanism of divergence has been gene duplications. While the *MHC* duplication predated metazoans, other smooth and striated-specific paralogs only diverged in bilaterians. Smooth and striated *MRLC* most likely arose by gene duplication in the bilaterian stem-line (Supplementary Figure 1). *Myosin essential light chain, actin* and *myocardin* paralogs split even later, in the vertebrate stem-line (Figure 7). Similarly, smooth and non-muscle *mhc* and *mrlc* paralogs only diverged in vertebrates. The *calponin-encoding* gene underwent parallel duplication and subfunctionalization in both annelids and chordates, giving rise to both specialized smooth muscle paralogs and more broadly expressed copies with a different domain structure (Figure 7—supplement 3). This slow and stepwise nature of the individuation process is consistent with studies showing that recently evolved paralogs can acquire differential expression between tissues that diverged long before in evolution (Force et al. 1999; Lan & Pritchard 2016).

Complementing gene duplication, the evolution and selective expression of entirely new apomeres also supported individuation: for example, Titin and all components of the Troponin complex are bilaterian novelties (Steinmetz et al. 2012). In vertebrates, the new gene *caldesmon* was incorporated in the smooth contractile module (Steinmetz et al. 2012).

### Smooth to striated myocyte conversion

Strikingly, visceral smooth myocytes were previously assumed to be a vertebrate innovation, as they are absent in fruit flies and nematodes (two groups which are in fact exceptions in this respect, at least from ultrastructural criteria (Figure 7—supplement 2A)). This view received apparent support from the fact that the vertebrate smooth and non-muscle myosin heavy chains (MHC) arose by vertebrate-specific duplication of a unique ancestral bilaterian gene, orthologous to *Drosophila* non-muscle MHC (Goodson & Spudich 1993) – which our results suggest reflects instead gradual individuation of pre-existing cell types (see above). Strikingly, the striated gut muscles of *Drosophila* closely resemble vertebrate and annelid smooth gut muscles by transcription factors *(nk3/bagpipe* (Azpiazu & Frasch 1993), *foxF/biniou* (Jakobsen et al. 2007a; Zaffran et al. 2001)), even though they express the fast/striated contractility module (Fyrberg et al. 1994; Fyrberg et al. 1990; Marín et al. 2004). If smooth gut muscles are ancestral for protostomes, as our results indicate, this suggests that the smooth contractile module was replaced by the fast/striated module in visceral myocytes during insect evolution. Interestingly, chromatin immunoprecipitation assays (Jakobsen et al. 2007a) show that the conserved visceral transcription factors *foxF/biniou* and *nk3/bagpipe* do not directly control contractility genes in *Drosophila* gut muscles (which are downstream *mef2* instead), but establish the morphogenesis and innervation of the visceral muscles, and control non-contractile effectors such as gap junctions – which are the properties these muscles seem to have conserved from their smooth ancestors. The striated gut myocytes of insects would thus represent a case of co-option of an effector module from another cell type, which happened at an unknown time during ecdysozoan evolution (Figure 7; Figure 7—supplement 1).

Another likely example of co-option is the vertebrate heart: vertebrate cardiomyocytes are striated and express fast myosin and troponin, but resemble smooth myocytes by developmental origin (from the splanchnopleura), function (automatic contraction and coupling by gap junctions) and terminal selector profile (Figure 1B). These similarities suggest that cardiomyocytes might stem from smooth myocytes that likewise co-opted the fast/striation module. Indicative of this possible ancestral state, the *Platynereis* dorsal pulsatile vessel (considered homologous to the vertebrate heart based on comparative anatomy and shared expression of *NK4/tinman* (Saudemont et al. 2008)) expresses the smooth, but not the striated, myosin heavy chain (Figure 3—supplement 3O-Q). An ancestral state reconstruction based on ultrastructural data further supports the notion that heart myocytes were smooth in the last common protostome/deuterostome ancestor, and independently acquired striation in at least 5 descendant lineages (Figure 7—supplement 2B) – usually in species with large body size and/or fast metabolism.

### Striated to smooth conversions

Smooth somatic muscles are occasionally found in bilaterians with slow or sessile lifestyles – for example in the snail foot (Faccioni-Heuser et al. 1999; Rogers 1969), the ascidian siphon (Meedel & Hastings 1993), and the sea cucumber body wall (Kawaguti & Ikemoto 1965). As an extreme (and isolated) example, flatworms lost striated muscles altogether, and their body wall musculature is entirely smooth (Rieger et al. 1991). Interestingly, in all cases that have been molecularly characterized, smooth somatic muscles express the same fast contractility module as their striated counterparts, including ST-MHC and the Troponin complex – in ascidians (Endo & Obinata 1981; Obinata et al. 1983), flatworms (Kobayashi et al. 1998; Witchley et al. 2013; Sulbarán et al. 2015), and the smooth myofibres of the bivalve catch muscle (Nyitray et al. 1994; Ojima & Nishita 1986). (It is unknown whether these also express *zasp* and *titin* in spite of the lack of striation). This suggests that these are somatic muscles having secondarily lost striation (in line with the sessile lifestyle of ascidians and bivalves, and with the complete loss of striated muscles in flatworms). Alternatively, they might represent remnants of ancestral smooth somatic fibres that would have coexisted alongside striated somatic fibres in the last common protostome/deuterostome ancestor. Interestingly, the fast contractile module is also expressed in acoel body wall smooth muscles (Chiodin et al. 2011); since acoels belong to a clade that might have branched off before all other bilaterians (Cannon et al. 2016) (but see (Philippe et al. 2011)), these could represent fast-contracting myocytes that never evolved striation in the first place, similar to those found in cnidarians. In all cases, the fast contractility module appears to represent a consistent synexpression group (i.e. its components are reliably expressed together), and a stable molecular profile of all bilaterian somatic muscles, regardless of the presence of morphologically overt striation. This confirms the notion that, even in cases of ambiguous morphology or ultrastructure, the molecular fingerprint of cell types holds clue to their evolutionary affinities.

### Implications for cell type evolution

In the above, genetically well-documented cases of cell type conversion (smooth to striated conversion in insect visceral myocytes and vertebrate cardiomyocytes), cells kept their ancestral CRC of terminal selector transcription factors, while changing the downstream effector modules. This supports the recent notion that CRCs confer an abstract identity to cell types, which remains stable in spite of turnover in downstream effectors (Wagner 2014) – just as *hox* genes impart conserved abstract identity to segments of vastly diverging morphologies (Deutsch 2005). Tracking cell type-specific CRCs through animal phylogeny thus represents a powerful means to decipher the evolution of cell types.

### Pre-bilaterian origins

If the existence of fast-contracting striated and slow-contracting smooth myocytes predated bilaterians – when and how did these cell types first split in evolution? The first evolutionary event that paved the way for the diversification of the smooth and striated contractility modules was the duplication of the striated myosin heavy chain-encoding gene into the striated isoform *ST-MHC* and the smooth/non-muscle isoform *SM-MHC*. This duplication occurred in single-celled ancestors of animals, before the divergence of filastereans and choanoflagellates (Steinmetz et al. 2012). Consistently, both *sm-mhc* and *st-mhc* are present in the genome of the filasterean *Ministeria* (though *st-mhc* was lost in other single-celled holozoans) (Sebé-Pedrós et al. 2014). Interestingly, *st-mhc* and *sm-mhc* expression appears to be segregated into distinct cell types in sponges, cnidarians (Steinmetz et al. 2012), and ctenophores (Dayraud et al. 2012), suggesting that a cell type split between slow and fast contractile cells is a common feature across early-branching metazoans (Figure 7). Given the possibility of *MHC* isoform co-option (as outlined above), it is yet unclear whether this split happened once or several times. The affinities of bilaterians and non-bilaterians contractile cells remain to be tested from data on the CRCs establishing contractile cell types in non-bilaterians.

## Conclusions

Our results indicate that the split between visceral smooth myocytes and somatic striated myocytes is the result of a long individuation process, initiated before the last common protostome/deuterostome ancestor. Fast- and slow-contracting cells expressing distinct variants of myosin II heavy chain *(ST-MHC* versus *SM-MHC*) acquired increasingly contrasted molecular profiles in a gradual fashion – and this divergence process continues to this day in individual bilaterian phyla. Blurring this picture of divergence, co-option events have led to the replacement of the slow contractile module by the fast one, leading to smooth-to-striated myocyte conversions. Our study showcases the power of molecular fingerprint comparisons centering on effector and selector genes to reconstruct cell type evolution (Arendt 2008). In the bifurcating phylogenetic tree of animal cell types (Liang et al. 2015), it remains an open question how the two types of contractile cells relate to other cell types, such as neurons (Mackie 1970) or cartilage (Lauri et al. 2014).

## Material and Methods

### Immunostainings and in situ hybridizations

Immunostaining, rhodamine-phalloidin staining, and simple and double WMISH were performed according to previously published protocols (Lauri et al. 2014). For all stainings not involving phalloidin, the following modifications: animals were mounted in 97% TDE/3% PTw for imaging following (Asadulina et al. 2012). Phalloidin-stained larvae were mounted in 1% DABCO/glycerol instead, as TDE was found to quickly disrupt phalloidin binding to F-actin. Confocal imaging of stained larvae was performed using a Leica SPE and a Leica SP8 microscope. Stacks were visualized and processed with ImageJ 1.49v. 3D renderings were performed with Imaris 8.1. Bright field Nomarski microscopy was performed on a Zeiss M1 microscope. Z-projections of Nomarski stacks were performed using Helicon Focus 6.7.1.

### Transmission electron microscopy

TEM was performed as previously published (Lauri et al. 2014).

### Pharmacological treatments

Brefeldin A was purchased from Sigma Aldrich (B7561) and dissolved in DMSO to a final concentration of 5 mg/mL. Animals were treated with 50 µg/mL Brefeldin A in 6-well plates filled with 5 mL filtered natural sea water (FNSW). Controls were treated with 1% DMSO (which is compatible with *Platynereis* development and survival without noticeable effect). Other neurotransmission inhibitors had no effect on *Platynereis* (as they elicited no impairment of locomotion): tetanus toxic (Sigma Aldrich T3194; 100 μg/mL stock in distilled water) up to 5 μg/mL; TTX (Latoxan, L8503; 1 mM stock) up to 10 μM; Myobloc (rimabotulinum toxin B; Solstice Neurosciences) up to 1%; saxitoxin 2 HCl (Sigma Aldrich NRCCRMSTXF) up to 1%; and neosaxitoxin HCl (Sigma Aldrich NRCCRMNEOC) up to 1%. (±)-2-Octanol was purchased from Sigma Aldrich and diluted to a final concentration of 2.5 mM (2 μL in 5 mL FNSW). (±)-2-Octanol treatment inhibited both locomotion and gut peristalsis, in line with the importance of gap junctions in motor neural circuits (Kawano et al. 2011; Kiehn & Tresch 2002). No sample size was computed before the experiments. At least 2 technical replicates were performed for each assay, with at least 5 biological replicates per sample per technical replicate. A technical replicate is a batch of treated individuals (together with their control siblings), and a biological replicate is a treated (or control sibling) individual.

### Live imaging of contractions

Animals were mounted in 3% low melting point agarose in FNSW (2-Hydroxyethylagarose, Sigma Aldrich A9414) between a slide and a cover slip (using 5 layers of adhesive tape for spacing) and imaged with a Leica SP8 confocal microscope. Fluorescent labeling of musculature was achieved either by microinjection of mRNAs encoding *GCaMP6s, LifeAct-EGFP* or *H2B-RFP*, or by incubation in 3 μM 0.1% FM-464FX (ThermoFisher Scientific, F34653). Contraction speed was calculated as (*l2-l1*)/(*l1*\*t), where *l1* is the initial length, *l2* the length after contraction, and *t* the duration of the contraction. Kymographs and wave speed quantifications were performed with the ImageJ Kymograph plugin: http://www.embl.de/eamnet/html/kymograph.html No sample size was computed before the experiments. At least 2 technical replicates were performed for each assay, with at least 2 biological replicates per sample per technical replicate. A technical replicate is a batch of treated individuals (together with their control siblings), and a biological replicate is a treated (or control sibling) individual.

### Ancestral state reconstruction

Ancestral state reconstructions were performed with Mesquite 3.04 using the Maximum Likelihood and Parsimony methods.

### Cloning

The following primers were used for cloning Platynereis genes using a mixed stages Platynereis cDNA library (obtained from 1, 2, 3, 5, 6, 10, and 14-days old larvae) and either the HotStart Taq Polymerase from Qiagen or the Phusion polymerase from New England BioLabs (for GC-rich primers):

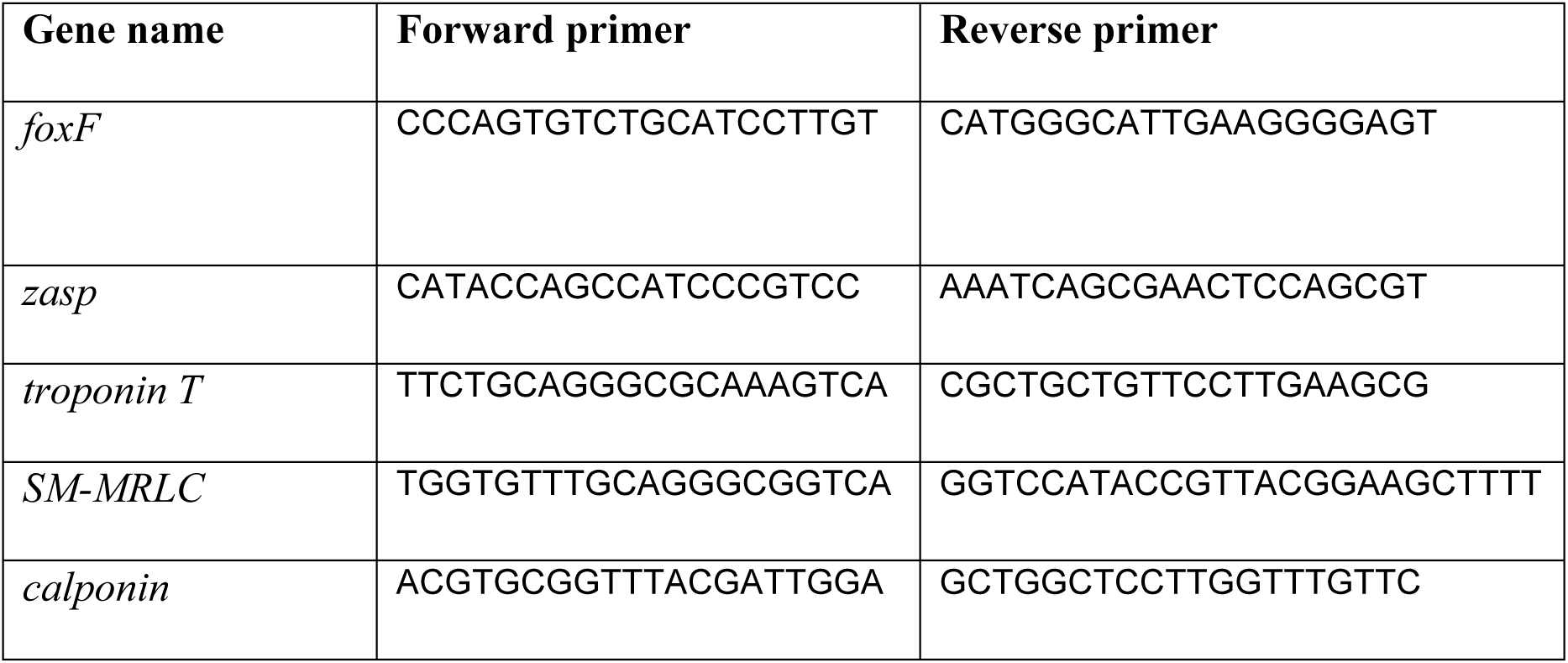
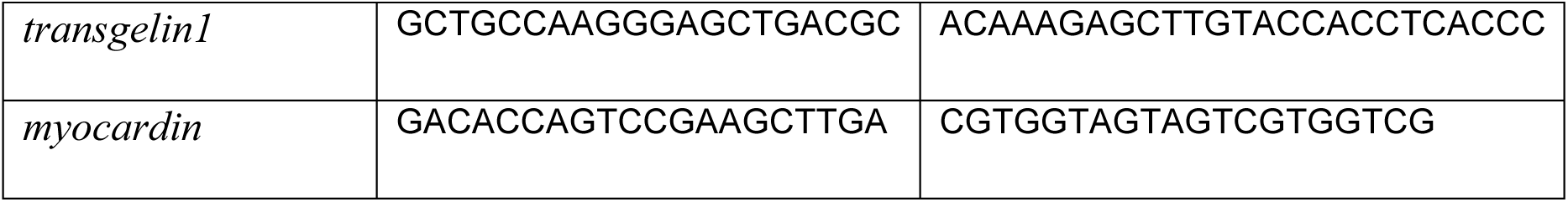

The following genes were retrieved from an EST plasmid stock: *SM-MHC* (as two independent clones that gave identical expression patterns) and *ST-MRLC*.

Other genes were previously published: *NK3* (Saudemont et al. 2008), *actin* and *ST-MHC* (under the name *mhc1-4*) (Lauri et al. 2014) and *GATA456* (Gillis et al. 2007).

## Acknowledgements

We thank Kaia Achim for the *hnf4* plasmid, Pedro Machado (Electron Microscopy Core Facility, EMBL Heidelberg) for embedding and sectioning samples for TEM, and the Arendt lab for feedback on the project. The work was supported by the European Research Council “Brain Evo-Devo” grant (TB, PB and DA), European Union’s Seventh Framework Programme project “Evolution of gene regulatory networks in animal development (EVONET)” [215781-2] (AL), the Zoonet EU-Marie Curie early training network [005624] (AF), the European Molecular Biology Laboratory (AL, AHLF, and PRHS) and the EMBL International PhD. Programme (TB, AHLF, PRHS, AL).

---

**Figure 2—figure supplement 1.**
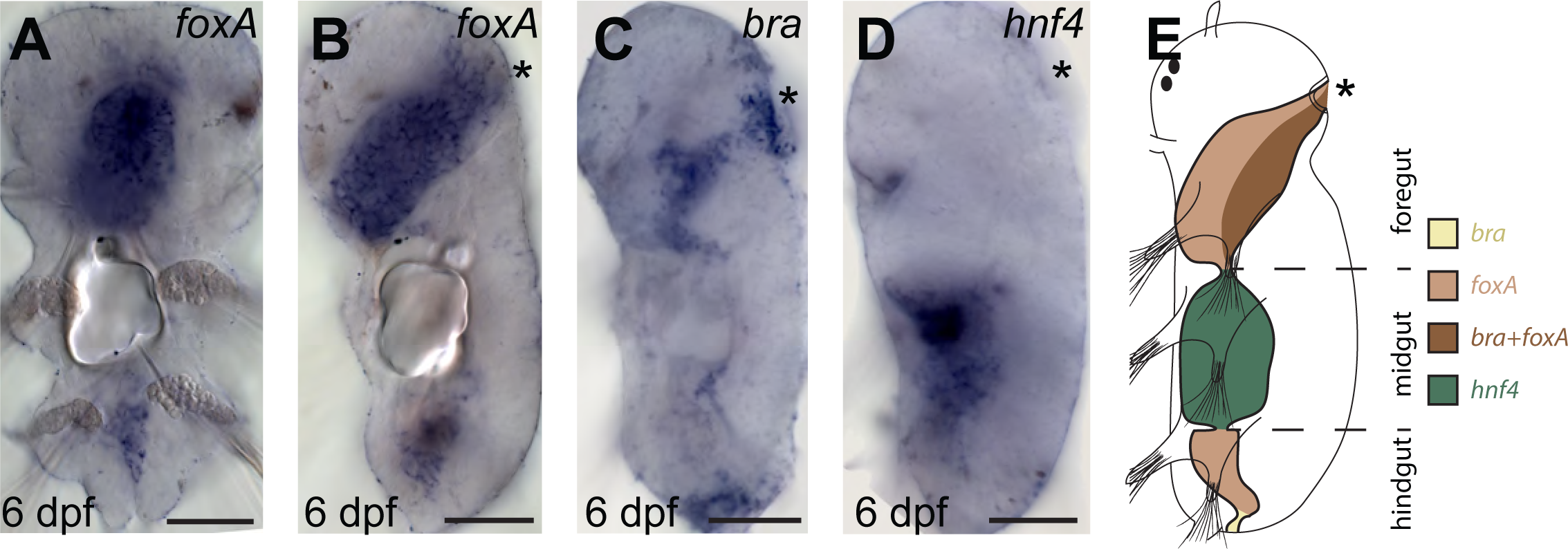
Gut patterning in *Platynereis* 6 dpf larvae. (A-D) WMISH for gut markers. (A) Ventral view, anterior is up. (B-D) Left lateral views, anterior is up, ventral is right. (E) Schematic of gut patterning in *Platynereis* late nectochaete larvae. Scale bar: 50 μm.

**Figure 3—supplement 1.**
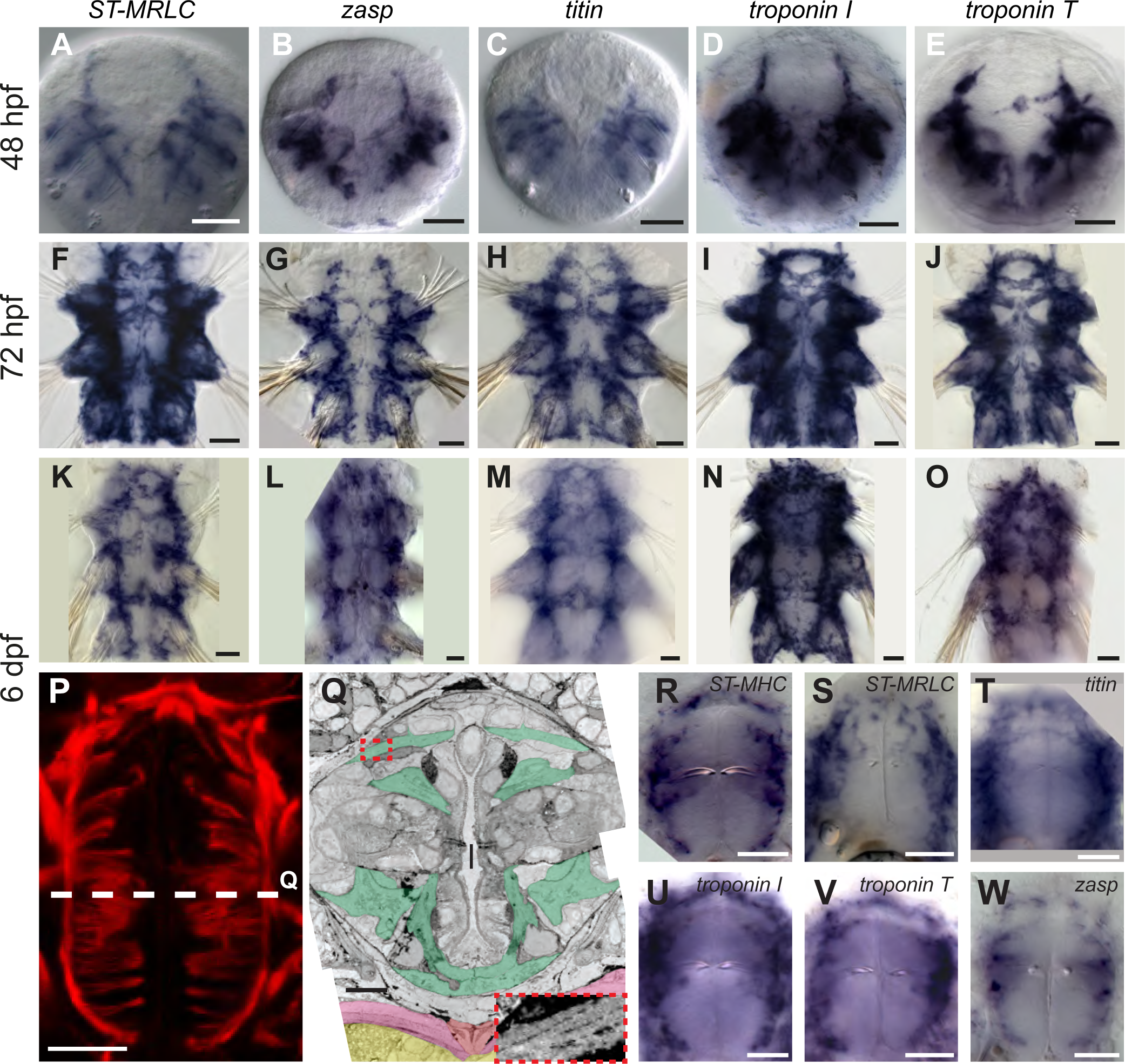
Expression of striated muscle markers in *Platynereis* larvae. (A-O) Larvae stained by WMISH and observed in bright field Nomarski microscopy. Ventral views, anterior side up. Scale bar is 20 µm for 48 hpf and 25 µm for the two other stages. (P) Foregut musculature visualized by rhodamine-phalloidin fluorescent staining. Z-projection of confocal planes. Ventral view, anterior side up. Scale bar: 20 µm. (Q) TEM micrograph of a cross-section of the foregut. Foregut muscles are colored green, axochord orange, ventral oblique muscles pink, ventral nerve cord yellow. Inset: zoom on the area in the red dashed box with enhanced contrast to visualize oblique striation. Scale bar: 10 µm. (R-W) WMISH for striated muscle markers expression in the foregut observed in Nomarski bright field microscopy, ventral views, anterior side up. Scale bar: 20 µm.

**Figure 3—supplement 2.**
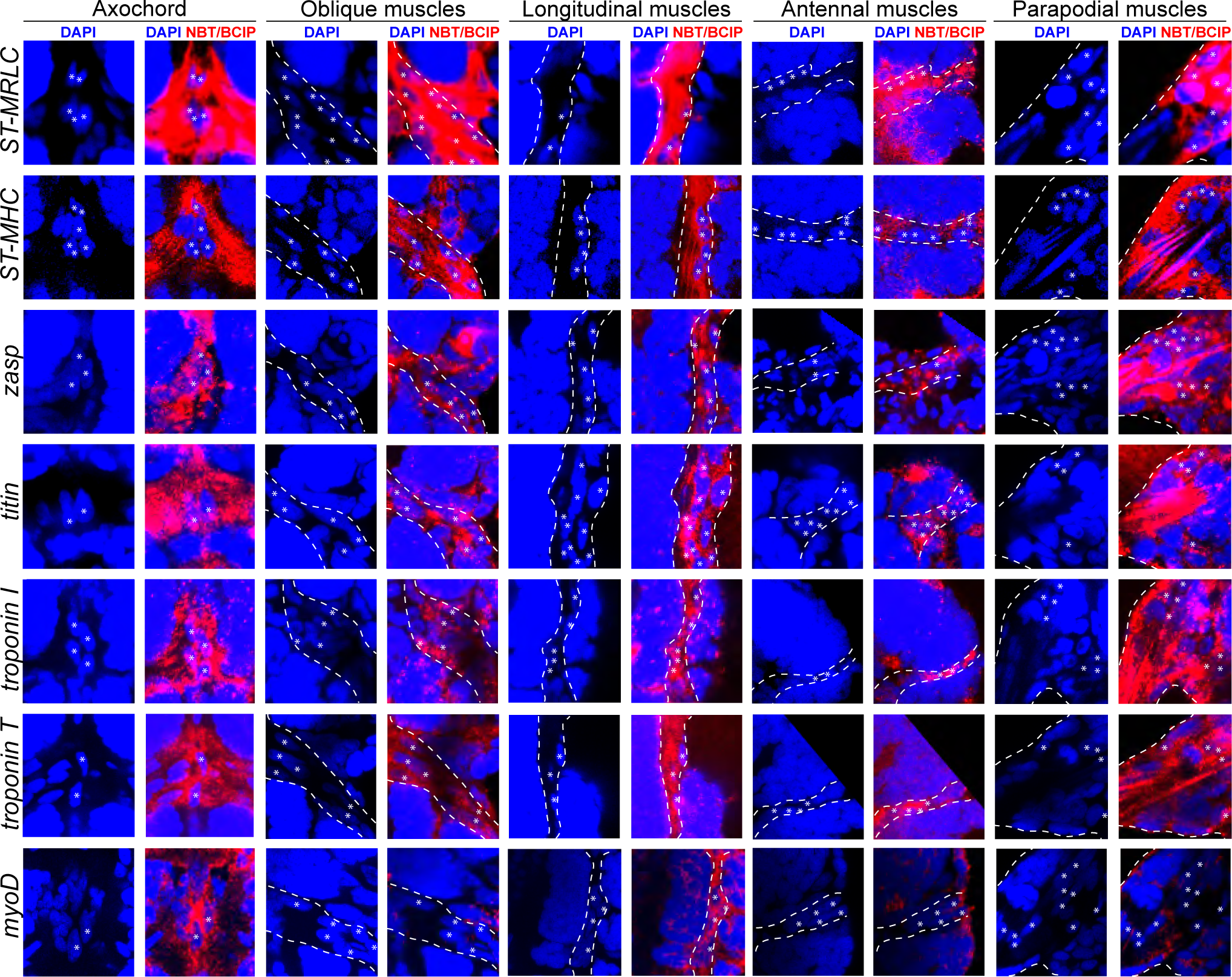
Expression of striated muscle markers in the 6 dpf *Platynereis* larva. Animals have been stained by WMISH and observed by confocal microscopy (DAPI fluorescence and NBT/BCIP 633 nm reflection). All striated effector genes are expressed in all somatic muscles examined. The transcription factor *myoD* is detectable in the axochord and in ventral longitudinal muscles, but not in other muscles.

**Figure 3—supplement 3.**
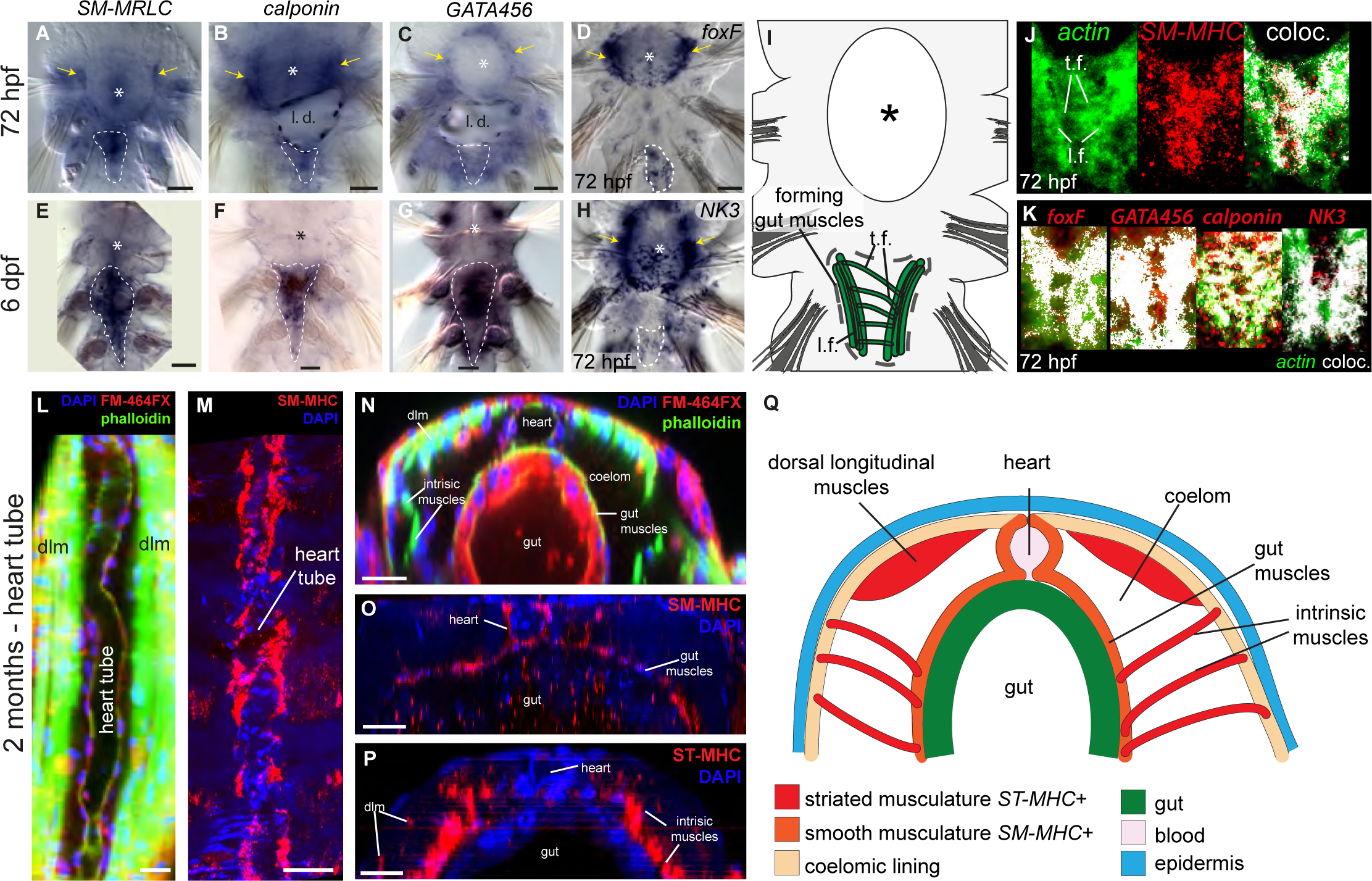
Expression of smooth muscle markers in *Platynereis* larvae. (A-H) Animals are stained by WMISH and observed by Nomarski bright field microscopy. Ventral views, anterior side up. Yellow arrows: expression in the foregut mesoderm. White dashed lines: outline of the midgut and hindgut (or their anlage at 3 dpf). Asterisk: stomodeum. (I) Schematic drawing of a 3 dpf larva (ventral view, anterior is up) showing the forming midgut and hindgut (dotted line) and the forming visceral myofibres in green (compared to double WMISH panels J and K). (J, K) Animals are stained by double WMISH and observed by confocal microscopy (cyanine 3 precipitate fluorescence for *actin*, NBT/BCIP precipitate reflection for other genes). Ventral views, anterior side up. Abbreviations: *t.f.:* transverse fibres; *l.f.:* longitudinal fibres. (L-Q) Molecular profile of the pulsatile dorsal vessel. All panels show 2 months-old juvenile worms. (L, M) Maximal Z-projections of confocal stacks. Dorsal view, anterior is up. (L) Dorsal musculature of a juvenile *Platynereis dumerilii* individual visualized by phalloidin-rhodamine (green) together with nuclear (DAPI, blue) and membrane (FM-464FX, red) stainings. The heart tube lies on the dorsal side, bordered by the somatic dorsal longitudinal muscles (*dlm*). (M) Expression of *SM-MHC* in the heart tube visualized by WMISH. (N) Virtual cross-section of the individual shown in A. Dorsal side up. Note the continuity of the muscular heart tube with gut musculature. (O) Virtual cross-section of the individual shown in B, showing continuous expression of *SM-MHC* in the heart and the midgut smooth musculature. Note the similarity to the *NK4/tinman* expression pattern documented in (Saudemont et al. 2008). (P) Virtual cross-section on an individual stained by WMISH for *ST-MHC* expression. Note the lack of expression in the heart, while expression is detected in intrinsic muscles that cross the internal cavity to suspend internal organs, and in dorsal longitudinal muscles (*dlm*). (Q) Schematic cross-section of a juvenile worm (dorsal side up) showing the shape, connections and molecular profile of the main muscle groups. Scale bar: 30 µm in all panels.

**Figure 3—supplement 4.**
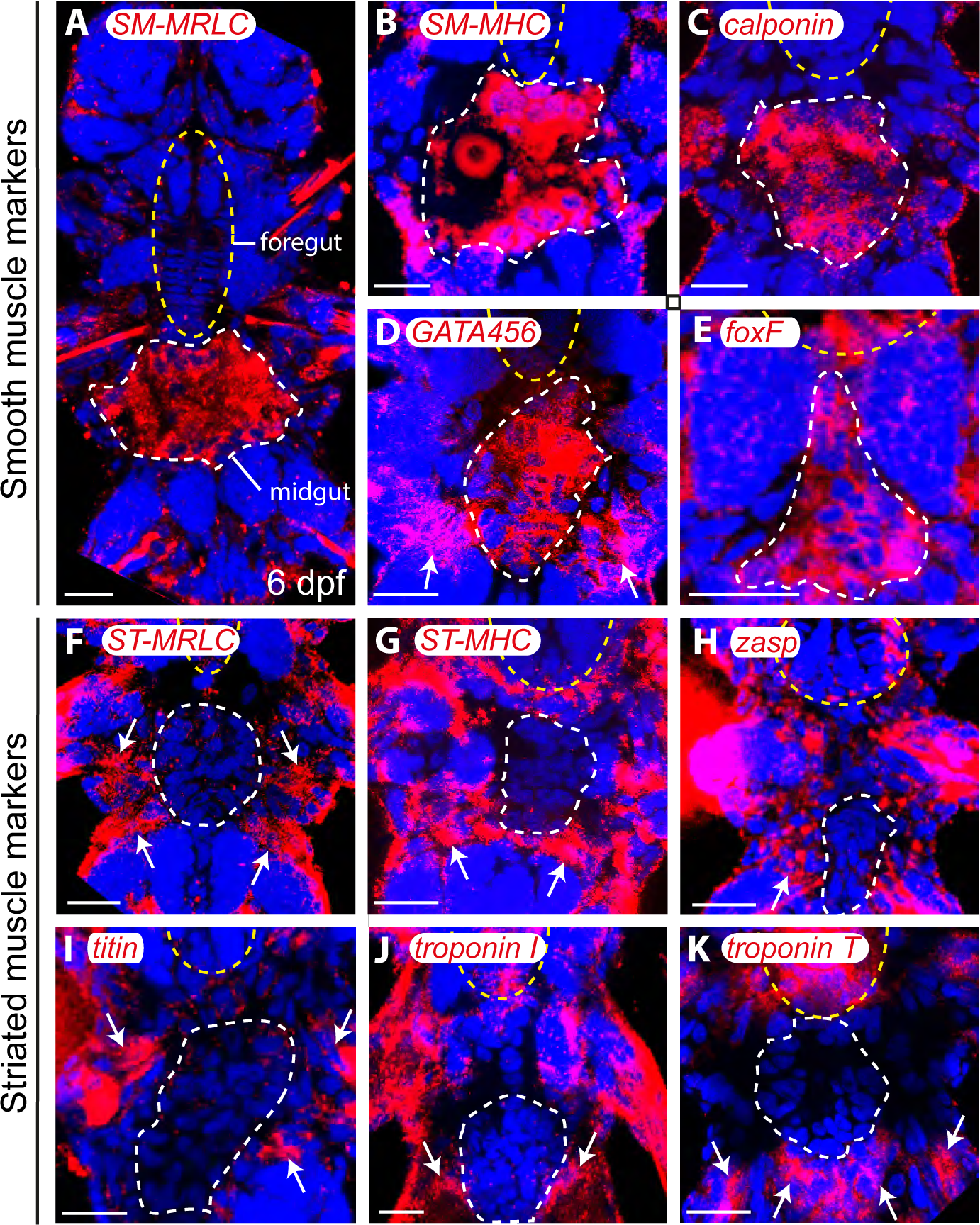
Molecular profile of midgut muscles in the 6 dpf larva. All panels are Z-projections of confocal planes, ventral views, anterior side up. Blue: DAPI, red: NBT/BCIP precipitate. White dashed line: midgut/hindgut, yellow dashed ellipse: stomodeum. (A-E) Smooth markers expression. White arrows indicate somatic expression of *GATA456* in the ventral oblique muscles. (F-K) Striated markers expression; none of them is detected in any gut cell. White arrows: somatic expression. Scale bar: 20 µm in all panels.

**Figure 3—supplement 5.**
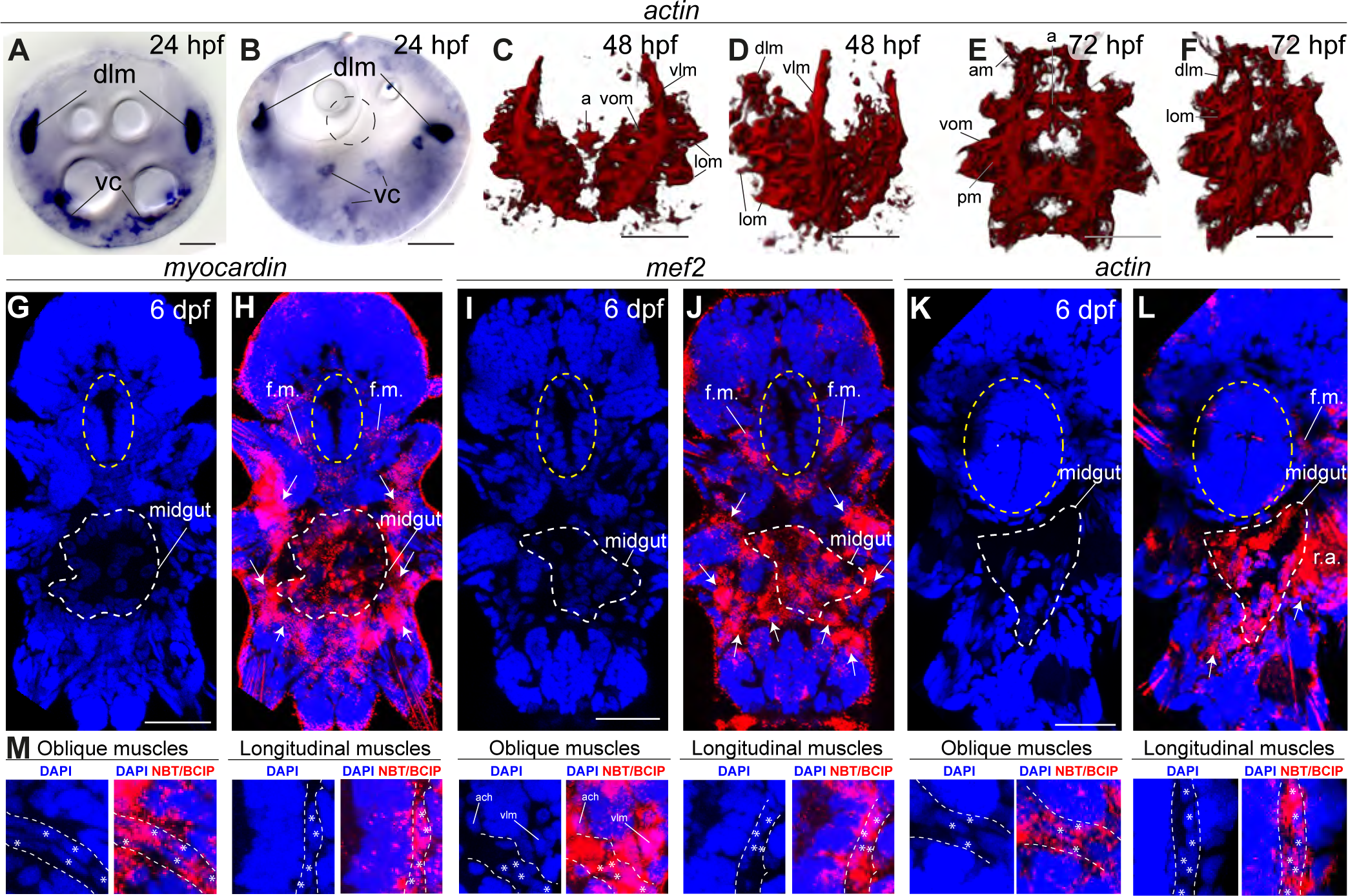
General muscle markers are expressed in both smooth and striated muscles. All panels show gene expression visualized by WMISH. (A-F) *actin* expression. (A-B) bright field micrographs in Nomarski optics. (A) is an apical view, (B) is a ventral view. Abbreviations: *dlm*, dorsal longitudinal muscles; vc, ventral mesodermal cells, likely representing future ventral musculature. (C-F) 3D rendering of confocal imaging of NBT/BCIP precipitate. (C, E) ventral views, anterior side up. (D, F) ventrolateral views, anterior side up. Abbreviations are as in Figure 1. (G-M) Z-projections of confocal stacks. Blue is DAPI, red is reflection signal of NBT/BCIP precipitate. (G-L) Ventral views, anterior side up. White dashed line: midgut, yellow ellipse: foregut. White arrows: somatic muscle expression. Abbreviations: *f.m.:* foregut muscles; *r.a.:* reflection artifact. (M) Expression in individual somatic muscles. Scale bar: 20 µm in all panels.

**Figure 7—supplement 1.**
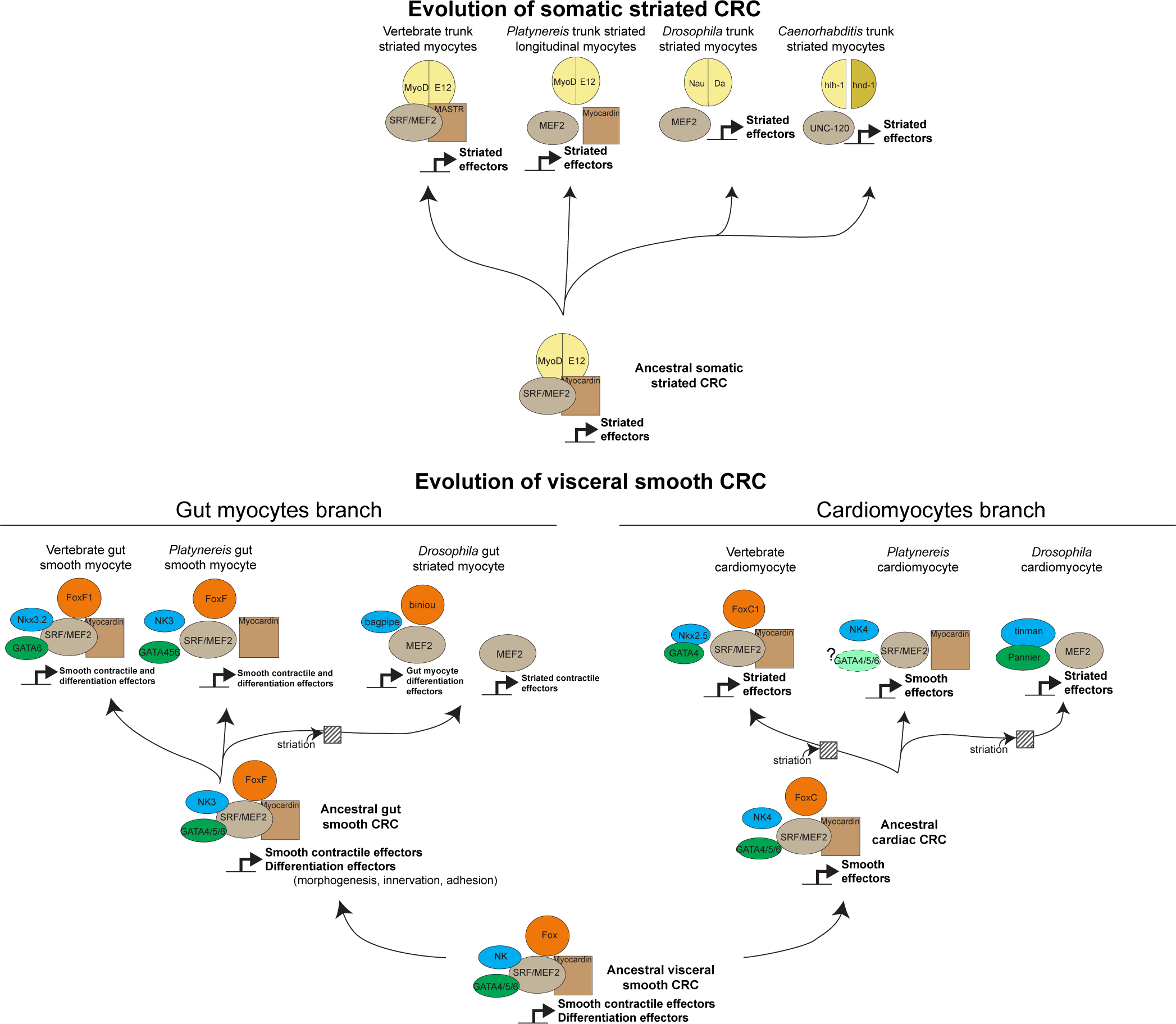
Evolution of myogenic Core Regulatory Complexes (CRC) in Bilateria. Transcription factor families are depicted as in Figure 1. Direct contact indicates proven binding. Co-option of the fast/striated module happened on three occasions: in *Drosophila* gut myocytes, and in cardiomyocytes of both vertebrates and *Drosophila*. Note that in both cases, composition of the CRC was maintained in spite of change in the effector module. In insect gut myocytes, replacement of the smooth by the striated module entailed a split of the CRC, with the ancient smooth CRC still controlling conserved differentiation genes (involved in adhesion, morphogenesis, axonal guidance, or formation of innexin gap junctions), while the striated contractile cassette is downstream Mef2 alone (Jakobsen et al. 2007b). It is less clear whether a similar split of CRC took place in striated cardiomyocytes.

**Figure 7—supplement 2.**
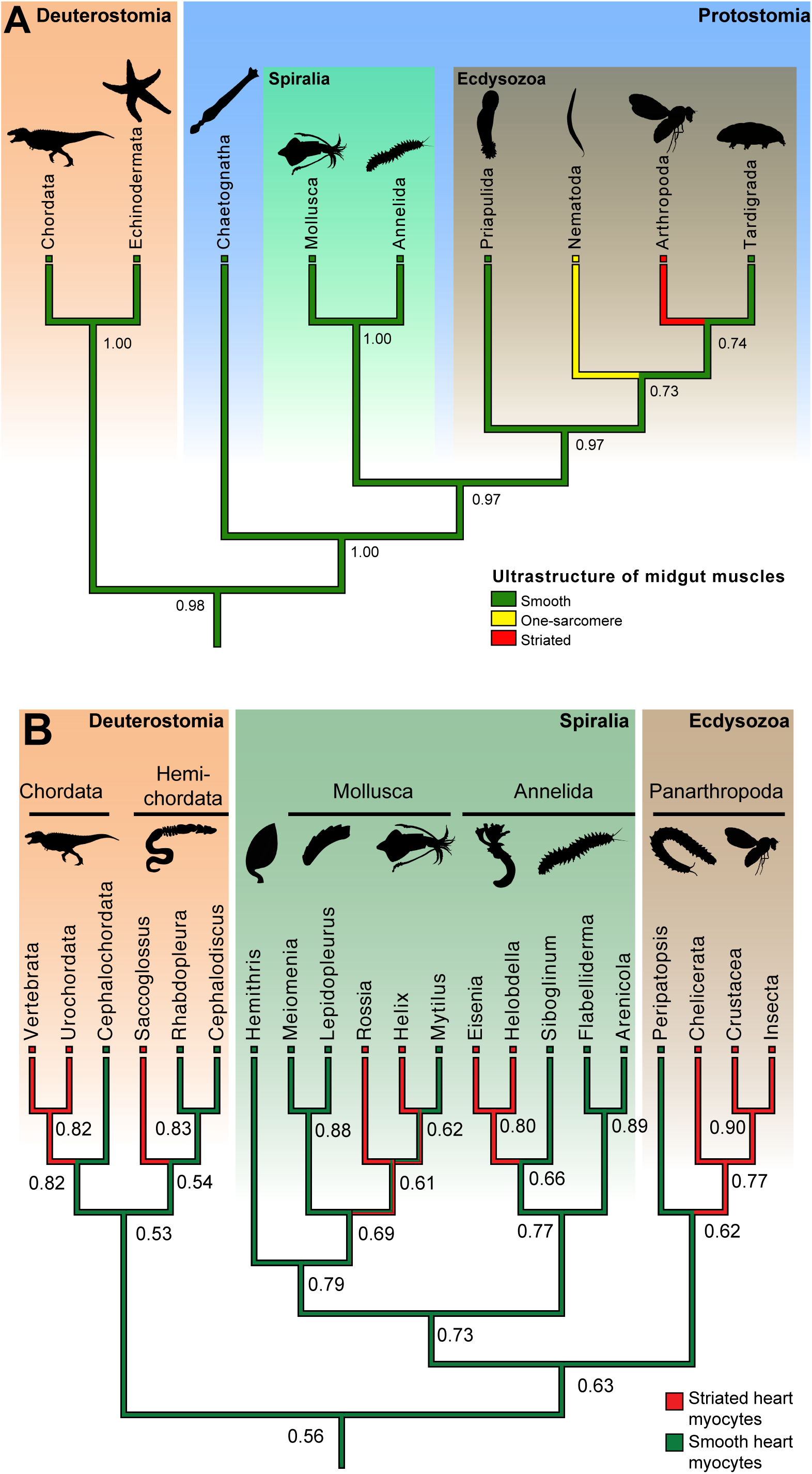
Ancestral state reconstructions of the ultrastructure of midgut/hindgut and heart myocytes. (A) Distribution, and ancestral state reconstruction, of midgut smooth muscles in Bilateria. Ancestral states were inferred using Parsimony and Maximum Likelihood (ML) (posterior probabilities indicated on nodes). Character states from: Chordata (Marieb & Hoehn 2015), Echinodermata (Feral & Massin 1982), Chaetognatha (Duvert & Salat 1995), Mollusca (Royuela et al. 2000), Annelida (Anderson & Ellis 1967), Priapulida (Carnevali & Ferraguti 1979), Nematoda (White 1988), Arthropoda (Goldstein & Burdette 1971), and Tardigrada (Shaw 1974). (B) Distribution, and ancestral state reconstruction, of cardiomyocyte ultrastructure in Bilateria. Ancestral states were inferred using Parsimony and ML (posterior probabilities indicated on nodes). Note that, due to the widespread presence of striated cardiomyocytes in bilaterians, the support value for an ancestral smooth ultrastructure in the ML method remain modest (0.56). This hypothesis receives independent support from the comparison of CRCs (Figure 1B, Figure 7—supplement 1) and developmental data (see Discussion). Character states follow the review by Martynova (Martynova 2004; Martynova 1995) and additional references for *Siboglinum* (Jensen & Myklebust 1975), chordates (Hirakow 1985), *Peripatopsis* (Nylund et al. 1988), arthropods (Tjønneland et al. 1987), *Meiomenia* (Reynolds et al. 1993), and *Lepidopleurus* #x00D8;kland 1980). In *Peripatopsis* and *Lepidopleurus*, some degree of alignment of dense bodies was detected (without being considered regular enough to constitute striation), suggesting these might represent intermediate configurations.

**Figure 7—supplement 3.**
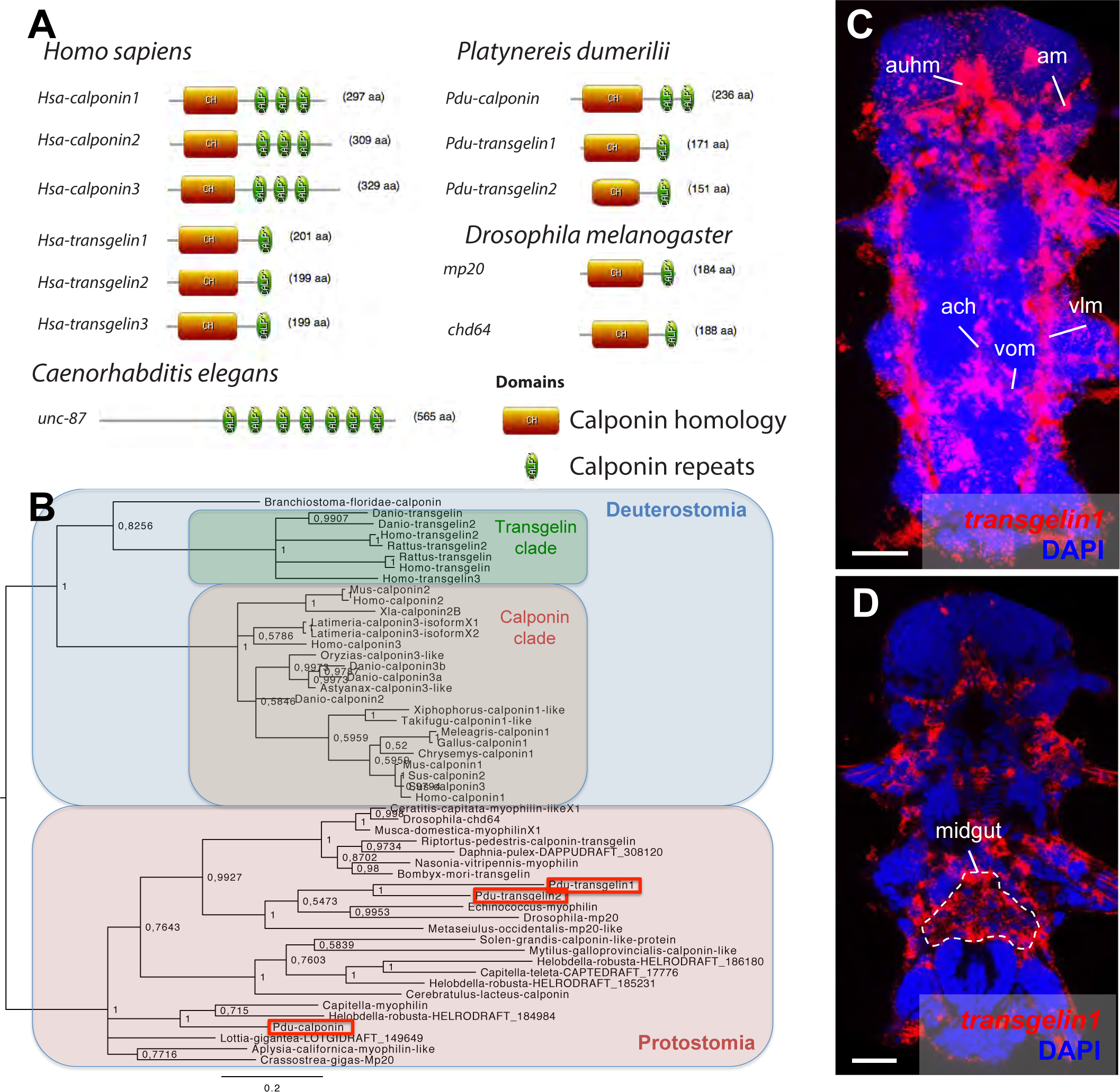
Domain structure, phylogeny and expression patterns of members of the *calponin* gene family. (A) Domain structure of calponin-related proteins in bilaterians. Calponin is characterized by a Calponin Homology (CH) domain with several calponin repeats, while Transgelin is characterized by a CH domain and a single calponin repeat. In vertebrates, Calponin proteins are specific smooth muscle markers, and the presence of multiple CH repeats allows them to stabilize actomyosin (Gimona et al. 2003). Transgelins, with a single calponin repeat, destabilize actomyosin (Gimona et al. 2003), and are expressed in other cell types such as podocytes (Gimona et al. 2003), lymphocytes (Francés et al. 2006), and striated muscles (transiently in mice (Li et al. 1996) and permanently in fruit flies (Ayme-Southgate et al. 1989)). (B) Maximum Likelihood phylogeny of the calponin/transgelin family based on alignment of the CH domain. Paralogs with calponin and transgelin structures evolved independently in vertebrates and *Platynereis* (Pdu, in red squares). (C) Expression patterns of *Pdu-transgelin1* Scale bar: 25 µm. As in vertebrates and insects, *transgelin1* is not smooth myocyte-specific, but also detected in striated myocytes.

**Supplementary Figure 1.**
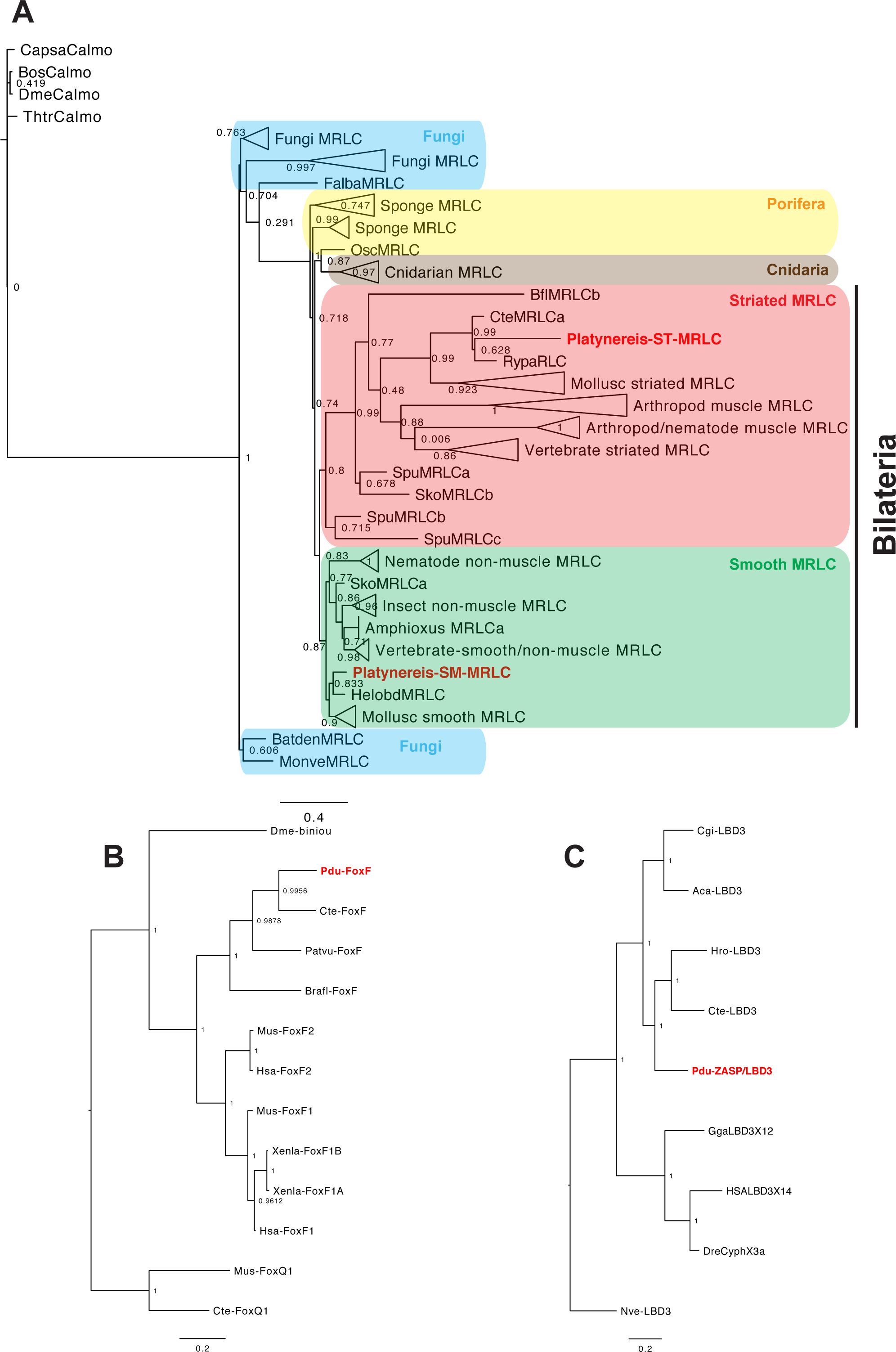
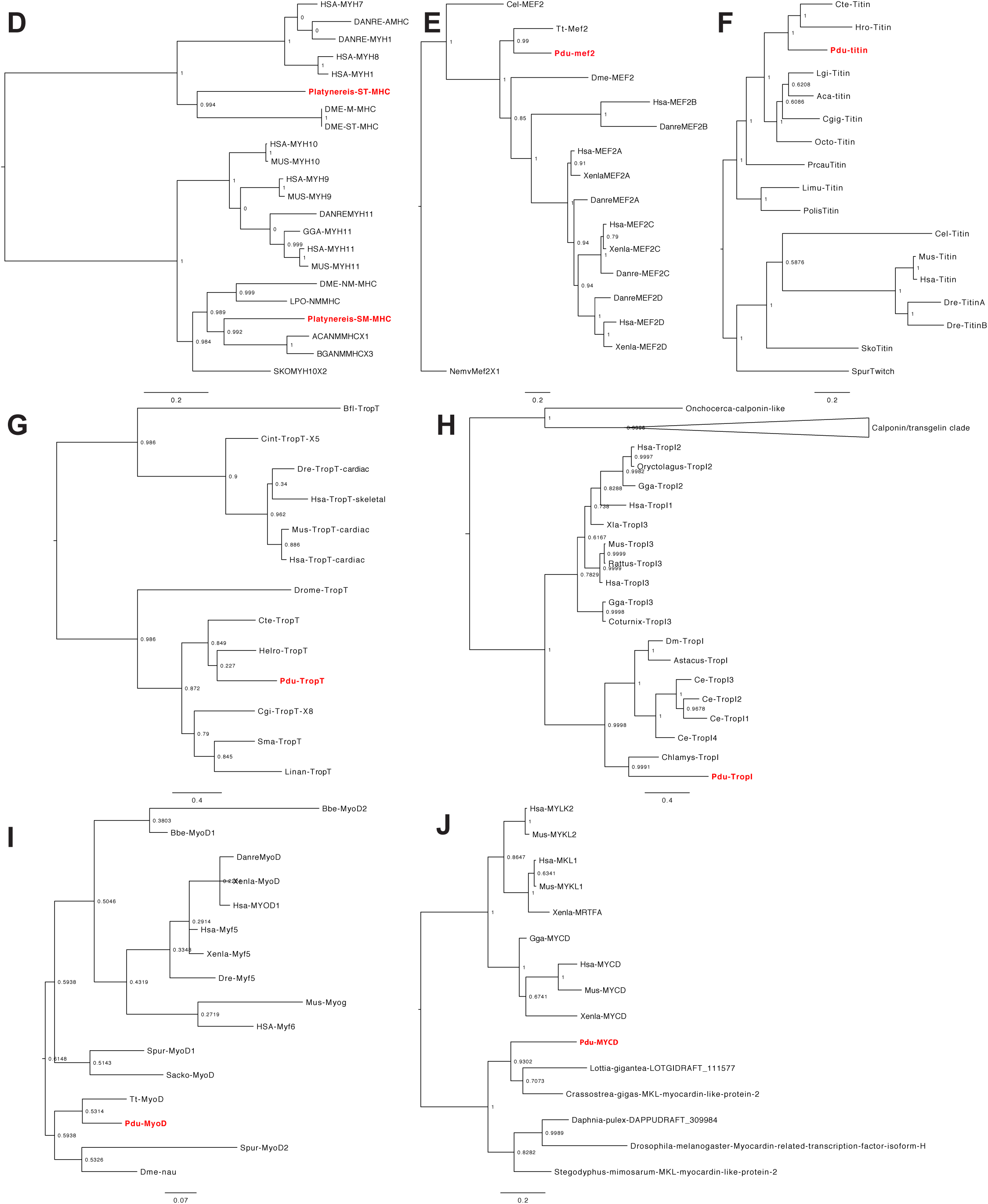
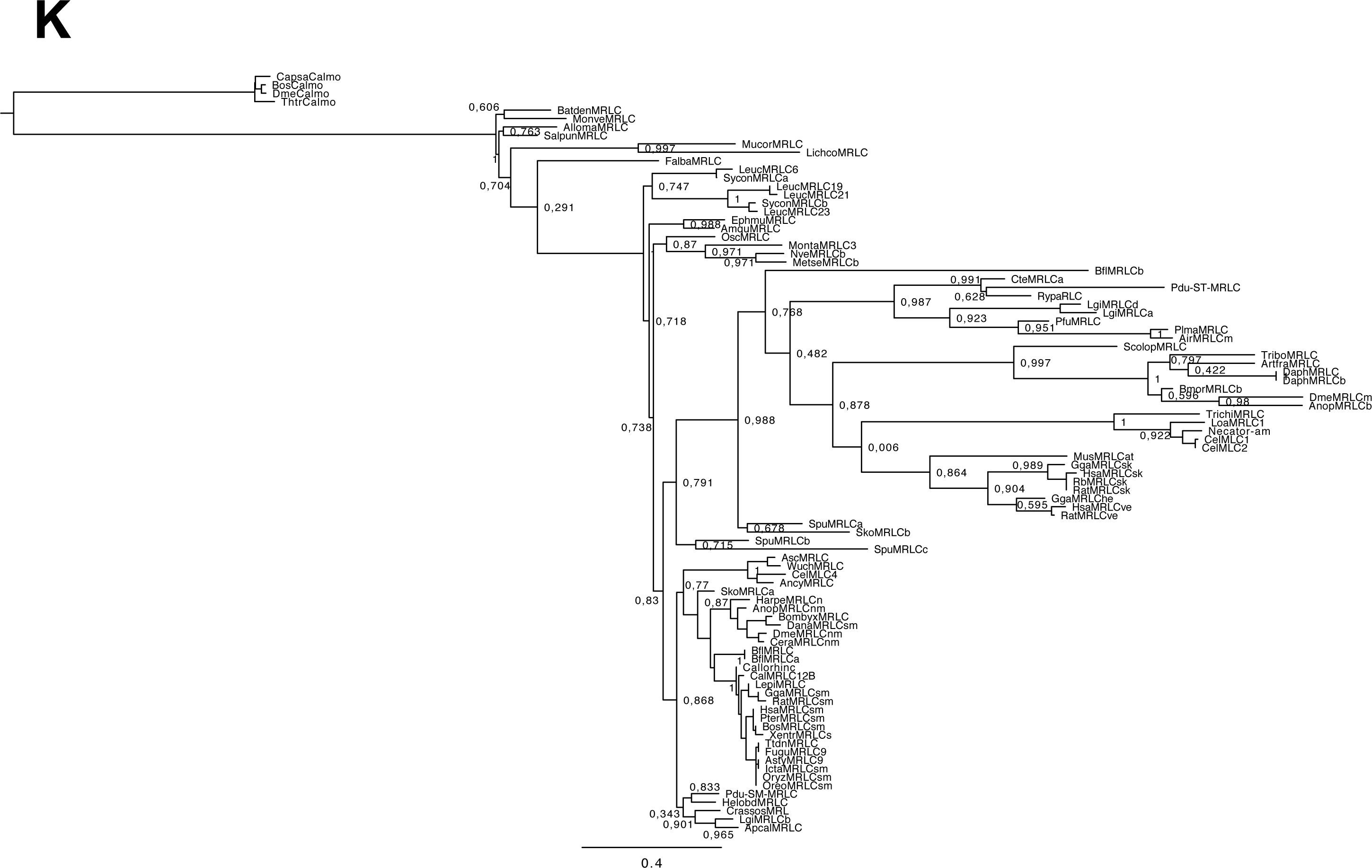
Phylogenetic trees of the markers investigated. (A) Simplified Maximum Likelihood (ML) tree for Myosin Regulatory Light Chain (full tree in panel M), rooted with Calmodulin, which shares an EF-hand calcium-binding domain with MRLC. (B) ML tree for FoxF, rooted with FoxQ1, the probable closest relative of the FoxF family (Shimeld et al. 2010). (C) MrBayes tree for bilaterian ZASP/LBD3, rooted with the cnidarian ortholog (Steinmetz et al. 2012). (D) ML tree for bilaterian Myosin Heavy Chain, rooted at the (pre-bilaterian) duplication between smooth and striated MHC (Steinmetz et al. 2012). (E) MrBayes tree for Mef2, rooted by the first splice isoform of the cnidarian ortholog (Genikhovich & Technau 2011). (F) MrBayes tree for Titin, rooted at the protostome/deuterostome bifurcation (Titin is a bilaterian novelty). (G) MrBayes tree for Troponin T, rooted at the protostome/deuterostome bifurcation (Troponin T is a bilaterian novelty). (H) MrBayes tree for Troponin I, rooted by the Calponin/Transgelin family, which shares an EF-hand calcium-binding domain with Troponin I. (I) MrBayes tree for MyoD, rooted at the protostome/deuterostome bifurcation (MyoD is a bilaterian novelty). (J) MrBayes tree for Myocardin, rooted at the protostome/deuterostome bifurcation (the *Drosophila myocardin* ortholog is established (Han et al. 2004)). (K) Complete MRLC tree. Species names abbreviations: Pdu: *Platynereis dumerilii*; Xenla: *Xenopus laevis;* Mus: *Mus musculus*; Hsa: *Homo sapiens*; Dre: *Danio rerio*; Gga: *Gallus gallus*; Dme: *Drosophila melanogaster;* Cte: *Capitella teleta;* Patvu: *Patella vulgata;* Brafl: *Branchiostoma floridae*; Nve or Nemv: *Nematostella vectensis;* Acdi: *Acropora digitifera*; Expal: *Exaiptasia pallida*; Rat: *Rattus norvegicus*; Sko: *Saccoglossus kowalevskii;* Limu or Lpo: *Limulus polyphemus*; Trib or Trca: *Tribolium castaneum*; Daph: *Daphnia pulex*; Prcau: *Priapulus caudatus;* Cgi or Cgig: *Crassostrea gigas;* Ling or Linan: *Lingula anatina*; Hdiv: *Haliotis diversicolor*; Apcal or Aca: *Aplysia californica*; Spu: *Strongylocentrotuspurpuratus*; Poli or Polis: *Polistes dominula*; Cin or Cint: *Ciona intestinalis*; Hro: *Helobdella robusta*; Bos: Bos *taurus*; Capsa: *Capsaspora owczarzaki*; Thtr: *Thecamonas trahens*; Lpo: ; Bga: *Biomphalaria glabrata;* Cel: *Caenorhabditis elegans;* Tt: *Terebratalia transversa;* Octo: *Octopus vulgaris;* Sma: *Schmidtea mediterranea;* Bbe: *Branchiostoma belcheri*, Batden: *Batrachochytrium dendrobaditis;* Monve: *Mortierella verticillata;* Alloma: *Allomyces macrogynus;* Salpun: *Spizellomyces punctatus;* Mucor: *Mucor racemosus;* Lichco: *Lichtheimia corymbifera;* Ephmu: *Ephydatia muelleri;* Sycon: *Sycon ciliatum;* Amqu: *Amphimedon queenslandica;* Osc: *Oscarella lobularis;* Metse: *Metridium senile;* Pfu: *Pinctada fucata;* Rypa: *Riftia pachyptila;* Plma: *Placopecten magellanicus;* Air: *Argopecten irradians;* Scolop: *Scolopendra gigantea;* Artfra: *Artemia franciscana;* Bmor: *Bombyx mori;* Loa: *Loa loa;* Necator-am: *Necator americanus;* Trichi: *Trichinella spiralis;* Asc: *Ascaris lumbricoides;* Wuch: *Wucheria bancrofti;* Ancy: *Ancylostoma duodenale;* Callorinc: *Callorhinchus milii;* Dana: *Danaus plexippus;* Anop: *Anopheles gambiae;* Asty: *Astyanax mexicanus;* Oreo: *Oreochromis niloticus;* Icta: *Ictalurus punctatus*.

## References

Alberts, B. et al., 2014. Molecular Biology of the Cell 6 edition., New York, NY: Garland Science.

Anderson, W.A. & Ellis, R.A., 1967. A comparative electron microscope study of visceral muscle fibers in Cambarus, Drosophila and Lumbricus. Zeitschrift für Zellforschung und Mikroskopische Anatomie, 79(4), pp.581–591.

Andrikou, C. et al., 2013. Myogenesis in the sea urchin embryo: the molecular fingerprint of the myoblast precursors. EvoDevo, 4(1), p.33.

Arendt, D., 2003. Evolution of eyes and photoreceptor cell types. International Journal of Developmental Biology, 47(7/8), pp.563–572.

Arendt, D. et al., 2016. Evolution of sister cell types by individuation. Nature Reviews Genetics, submitted.

Arendt, D., 2008. The evolution of cell types in animals: emerging principles from molecular studies. Nature Reviews. Genetics, 9(11), pp.868–882.

Asadulina, A. et al., 2012. Whole-body gene expression pattern registration in Platynereis larvae. EvoDevo, 3, p.27.

Au, Y. et al., 2004. Solution Structure of ZASP PDZ Domain: Implications for Sarcomere Ultrastructure and Enigma Family Redundancy. Structure, 12(4), pp.611–622.

Ayme-Southgate, A. et al., 1989. Characterization of the gene for mp20: a Drosophila muscle protein that is not found in asynchronous oscillatory flight muscle. The Journal of cell biology, 108(2), pp.521–531.

Azpiazu, N. & Frasch, M., 1993. tinman and bagpipe: two homeo box genes that determine cell fates in the dorsal mesoderm of Drosophila. Genes & Development, 7(7b), pp.1325–1340.

Bárány, M., 1967. ATPase activity of myosin correlated with speed of muscle shortening. The Journal of general physiology, 50(6), pp.197–218.

Beall, C.J. & Fyrberg, E., 1991. Muscle abnormalities in Drosophila melanogaster heldup mutants are caused by missing or aberrant troponin-I isoforms. The Journal of Cell Biology, 114(5), pp.941–951.

Black, B.L. & Olson, E.N., 1998. Transcriptional control of muscle development by myocyte enhancer factor-2 (MEF2) proteins. Annual Review of Cell and Developmental Biology, 14, pp.167–196.

Blais, A. et al., 2005. An initial blueprint for myogenic differentiation. Genes & Development, 19(5), pp.553–569.

Bohrmann, J. & Haas-Assenbaum, A., 1993. Gap junctions in ovarian follicles of Drosophila melanogaster: inhibition and promotion of dye-coupling between oocyte and follicle cells. Cell and Tissue Research, 273(1), pp.163–173.

Brunet, T. & Arendt, D., 2016. From damage response to action potentials: early evolution of neural and contractile modules in stem eukaryotes. Philosophical Transactions of the Royal Society of London. Series B, Biological Sciences, 371(1685), p.20150043.

Bulbring, E. & Crema, A., 1959. The release of 5-hydroxytryptamine in relation to pressure exerted on the intestinal mucosa. The Journal of Physiology, 146(1), pp.18–28.

Burke, R.D., 1981. Structure of the digestive tract of the pluteus larva of Dendraster excentricus (Echinodermata: Echinoida). Zoomorphology, 98(3), pp.209–225.

Burr, A. & Gans, C., 1998. Mechanical Significance of Obliquely Striated Architecture in Nematode Muscle. The Biological Bulletin, 194(1), pp.1–6.

Cannon, J.T. et al., 2016. Xenacoelomorpha is the sister group to Nephrozoa. Nature, 530(7588), pp.89–93.

Carnevali, M.D.C. & Ferraguti, M., 1979. Structure and ultrastructure of muscles in the priapulid Halicryptus spinulosus: functional and phylogenetic remarks. Journal of the Marine Biological Association of the United Kingdom, 59(3), pp.737–748.

Carson, J.A., Schwartz, R.J. & Booth, F.W., 1996. SRF and TEF-1 control of chicken skeletal alpha-actin gene during slow-muscle hypertrophy. American Journal of Physiology – Cell Physiology, 270(6), pp.C1624–C1633.

Chiodin, M. et al., 2011. Molecular architecture of muscles in an acoel and its evolutionary implications. Journal of Experimental Zoology Part B: Molecular and Developmental Evolution, 316B(6), pp.427–439.

Corsi, A.K. et al., 2000. Caenorhabditis elegans twist plays an essential role in non-striated muscle development. Development, 127(10), pp.2041–2051.

Creemers, E.E. et al., 2006. Coactivation of MEF2 by the SAP Domain Proteins Myocardin and MASTR. Molecular Cell, 23(1), pp.83–96.

Dayraud, C. et al., 2012. Independent specialisation of myosin II paralogues in muscle vs. non-muscle functions during early animal evolution: a ctenophore perspective. BMC evolutionary biology, 12(1), p.107.

Denes, A.S. et al., 2007. Molecular architecture of annelid nerve cord supports common origin of nervous system centralization in bilateria. Cell, 129(2), pp.277–288.

Deutsch, J., 2005. Hox and wings. BioEssays: News and Reviews in Molecular, Cellular and Developmental Biology, 27(7), pp.673–675.

Durocher, D. et al., 1997. The cardiac transcription factors Nkx2–5 and GATA-4 are mutual cofactors. The EMBO journal, 16(18), pp.5687–5696.

Duvert, M. & Salat, C., 1995. Ultrastructural Studies of the Visceral Muscles of Chaetognaths. Acta Zoologica, 76(1), pp.75–87.

Endo, T. & Obinata, T., 1981. Troponin and Its Components from Ascidian Smooth Muscle. Journal of Biochemistry, 89(5), pp.1599–1608.

Faccioni-Heuser, M.C. et al., 1999. The pedal muscle of the land snail Megalobulimus oblongus (Gastropoda, Pulmonata): an ultrastructure approach. Acta Zoologica, 80(4), pp.325–337.

Faussone – Pellegrini, M.-S. & Thuneberg, L., 1999. Guide to the identification of interstitial cells of Cajal. Microscopy research and technique, 47(4), pp.248–266.

Feral, J.-P. & Massin, C., 1982. Digestive systems: Holothuroidea. Echinoderm nutrition, pp.191–212.

Finkbeiner, S., 1992. Calcium waves in astrocytes-filling in the gaps. Neuron, 8(6), pp.1101–1108.

Fischer, A.H., Henrich, T. & Arendt, D., 2010. The normal development of Platynereis dumerilii (Nereididae, Annelida). Frontiers in zoology, 7(1), p.1.

Force, A. et al., 1999. Preservation of Duplicate Genes by Complementary, Degenerative Mutations. Genetics, 151(4), pp.1531–1545.

Francés, R. et al., 2006. B-1 cells express transgelin 2: unexpected lymphocyte expression of a smooth muscle protein identified by proteomic analysis of peritoneal B-1 cells. Molecular immunology, 43(13), pp.2124–2129.

Fyrberg, C. et al., 1994. Drosophila melanogaster genes encoding three troponin-C isoforms and a calmodulin-related protein. Biochemical Genetics, 32(3–4), pp. 119—135.

Fyrberg, E. et al., 1990. Drosophila melanogaster troponin-T mutations engender three distinct syndromes of myofibrillar abnormalities. Journal of Molecular Biology, 216(3), pp.657–675.

Genikhovich, G. & Technau, U., 2011. Complex functions of Mef2 splice variants in the differentiation of endoderm and of a neuronal cell type in a sea anemone. Development, 138(22), pp.4911–4919.

Gho, M., 1994. Voltage-clamp analysis of gap junctions between embryonic muscles in Drosophila. The Journal of Physiology, 481(Pt 2), pp.371–383.

Gillis, W.J., Bowerman, B. & Schneider, S.Q., 2007. Ectoderm-and endomesoderm-specific GATA transcription factors in the marine annelid Platynereis dumerilli. Evolution & development, 9(1), pp.39–50.

Gimona, M. et al., 2003. Calponin repeats regulate actin filament stability and formation of podosomes in smooth muscle cells. Molecular biology of the cell, 14(6), pp.2482–2491.

Goldstein, M.A. & Burdette, W.J., 1971. Striated visceral muscle of Drosophila melanogaster. Journal of Morphology, 134(3), pp.315–334.

Goodson, H.V. & Spudich, J.A., 1993. Molecular evolution of the myosin family: relationships derived from comparisons of amino acid sequences. Proceedings of the National Academy of Sciences of the United States of America, 90(2), pp.659–663.

Gopalakrishnan, S. et al., 2015. A cranial mesoderm origin for esophagus striated muscles. Developmental cell, 34(6), pp.694–704.

Han, Z. et al., 2004. A myocardin-related transcription factor regulates activity of serum response factor in Drosophila. Proceedings of the National Academy of Sciences of the United States of America, 101(34), pp.12567–12572.

Hirakow, R., 1985. The vertebrate heart in relation to the protochordates. Fortschritt der Zoologie, 30, pp.367–369.

Hobert, O., 2016. Chapter Twenty-Five – Terminal Selectors of Neuronal Identity. In P. M. Wassarman, ed. Current Topics in Developmental Biology. Essays on Developmental Biology, Part A. Academic Press, pp. 455–475. Available at: http://www.sciencedirect.com/science/article/pii/S0070215315002148 [Accessed June 23, 2016].

Hoggatt, A.M. et al., 2013. The Transcription Factor Foxf1 Binds to Serum Response Factor and Myocardin to Regulate Gene Transcription in Visceral Smooth Muscle Cells. Journal of Biological Chemistry, 288(40), pp.28477–28487.

Hooper, S.L., Hobbs, K.H. & Thuma, J.B., 2008. Invertebrate muscles: thin and thick filament structure; molecular basis of contraction and its regulation, catch and asynchronous muscle. Progress in neurobiology, 86(2), pp.72–127.

Hooper, S.L. & Thuma, J.B., 2005. Invertebrate muscles: muscle specific genes and proteins. Physiological reviews, 85(3), pp.1001–1060.

Jakobsen, J.S. et al., 2007a. Temporal ChIP-on-chip reveals Biniou as a universal regulator of the visceral muscle transcriptional network. Genes & Development, 21(19), pp.2448–2460.

Jakobsen, J.S. et al., 2007b. Temporal ChIP-on-chip reveals Biniou as a universal regulator of the visceral muscle transcriptional network. Genes & Development, 21(19), pp.2448–2460.

Jensen, H., 1974. Ultrastructural studies of the hearts in Arenicola marina L. (Annelida: Polychaeta). Cell and Tissue Research, 156(1), pp.127–144.

Jensen, H. & Myklebust, R., 1975. Ultrastructure of muscle cells in Siboglinum fiordicum (Pogonophora). Cell and tissue research, 163(2), pp.185–197.

Kamm, K.E. & Stull, J.T., 1985. The function of myosin and myosin light chain kinase phosphorylation in smooth muscle. Annual review of pharmacology and toxicology, 25(1), pp.593–620.

Katzemich, A. et al., 2011. Muscle type-specific expression of Zasp52 isoforms in Drosophila. Gene Expression Patterns, 11(8), pp.484–490.

Kawaguti, S. & Ikemoto, N., 1965. Electron microscopy on the longitudinal muscle of the sea-cucumber (Stichopus japonicus). S. Ebashi et al., Edits. Molecular biology of muscular contraction. Tokyo: Igaku Shoin Ltd. XII, pp.124–131.

Kawano, T. et al., 2011. An imbalancing act: gap junctions reduce the backward motor circuit activity to bias C. elegans for forward locomotion. Neuron, 72(4), pp.572–586.

Kiehn, O. & Tresch, M.C., 2002. Gap junctions and motor behavior. Trends in Neurosciences, 25(2), pp. 108–115.

Kierszenbaum, A.L. & Tres, L., 2015. Histology and Cell Biology: An Introduction to Pathology, 4e 4 edition., Philadelphia, PA: Saunders.

Kobayashi, C. et al., 1998. Identification of Two Distinct Muscles in the Planarian Dugesia japonica by their Expression of Myosin Heavy Chain Genes. Zoological Science, 15(6), pp.861–869.

Labeit, S. & Kolmerer, B., 1995. Titins: Giant Proteins in Charge of Muscle Ultrastructure and Elasticity. Science, 270(5234), pp.293–296.

Lan, X. & Pritchard, J.K., 2016. Coregulation of tandem duplicate genes slows evolution of subfunctionalization in mammals. Science, 352(6288), pp.1009–1013.

Lauri, A. et al., 2014. Development of the annelid axochord: insights into notochord evolution. Science (New York, N.Y.), 345(6202), pp.1365–1368.

Lee, H.-H., Zaffran, S. & Frasch, M., 2006. Development of the larval visceral musculature. In Muscle Development in Drosophila. Springer, pp. 62–78.

Lee, Y. et al., 1998. The Cardiac Tissue-Restricted Homeobox Protein Csx/Nkx2.5 Physically Associates with the Zinc Finger Protein GATA4 and Cooperatively Activates Atrial Natriuretic Factor Gene Expression. Molecular and Cellular Biology, 18(6), pp.3120–3129.

Leininger, S. et al., 2014. Developmental gene expression provides clues to relationships between sponge and eumetazoan body plans. Nature Communications, 5, p.3905.

Li, L. et al., 1996. SM22a, a Marker of Adult Smooth Muscle, Is Expressed in Multiple Myogenic Lineages During Embryogenesis. Circulation Research, 78(2), pp.188–195.

Liang, C. et al., 2015. The statistical geometry of transcriptome divergence in cell-type evolution and cancer. Nature Communications, 6, p.6066.

Mackie, G.O., 1970. Neuroid conduction and the evolution of conducting tissues. The Quarterly Review of Biology, 45(4), pp.319–332.

Malo, M. et al., 2000. Effect of brefeldin A on acetylcholine release from glioma C6BU-1 cells. Neuropharmacology, 39(11), pp.2214–2221.

Marieb, E.N. & Hoehn, K.N., 2015. Human Anatomy & Physiology 10 edition., Boston: Pearson.

Marín, M.-C., Rodríguez, J.-R. & Ferrús, A., 2004. Transcription of Drosophila Troponin I Gene Is Regulated by Two Conserved, Functionally Identical, Synergistic Elements. Molecular Biology of the Cell, 15(3), pp.1185–1196.

Martindale, M.Q., Pang, K. & Finnerty, J.R., 2004. Investigating the origins of triploblasty:mesodermal’gene expression in a diploblastic animal, the sea anemone Nematostella vectensis (phylum, Cnidaria; class, Anthozoa). Development, 131(10), pp.2463–2474.

Martín-Durán, J.M. & Hejnol, A., 2015. The study of Priapulus caudatus reveals conserved molecular patterning underlying different gut morphogenesis in the Ecdysozoa. BMC biology, 13, p.29.

Martynova, M.G., 1995. Possible Cellular Mechanisms of Heart Muscle Growth in Invertebratea. Annals of the New York Academy of Sciences, 752(1), pp.149–157.

Martynova, M.G., 2004. Proliferation and Differentiation Processes in the Heart Muscle Elements in Different Phylogenetic Groups. In B.-I. R. of Cytology, ed. Academic Press, pp. 215–250. Available at: http://www.sciencedirect.com/science/article/pii/S0074769604350059 [Accessed June 17, 2016].

McKeown, C.R., Han, H.-F. & Beckerle, M.C., 2006. Molecular characterization of the Caenorhabditis elegans ALP/Enigma gene alp-1. Developmental Dynamics, 235(2), pp.530–538.

Meadows, S.M. et al., 2008. The myocardin-related transcription factor, MASTR, cooperates with MyoD to activate skeletal muscle gene expression. Proceedings of the National Academy of Sciences, 105(5), pp.1545–1550.

Meedel, T.H. & Hastings, K.E., 1993. Striated muscle-type tropomyosin in a chordate smooth muscle, ascidian body-wall muscle. Journal of Biological Chemistry, 268(9), pp.6755–6764.

Mill, P.J. & Knapp, M.F., 1970. The Fine Structure of Obliquely Striated Body Wall Muscles in the Earthworm, Lumbricus Terrestris LINN. Journal of Cell Science, 7(1), pp.233–261.

Misumi, Y. et al., 1986. Novel blockade by brefeldin A of intracellular transport of secretory proteins in cultured rat hepatocytes. Journal of Biological Chemistry, 261(24), pp.11398–11403.

Molkentin, J.D. et al., 1995. Cooperative activation of muscle gene expression by MEF2 and myogenic bHLH proteins. Cell, 83(7), pp.1125–1136.

Morin, S. et al., 2000. GATA-dependent recruitment of MEF2 proteins to target promoters. The EMBO journal, 19(9), pp.2046–2055.

Musser, J.M. & Wagner, G.P., 2015. Character trees from transcriptome data: Origin and individuation of morphological characters and the so-called “species signal.” Journal of Experimental Zoology. Part B, Molecular and Developmental Evolution, 324(7), pp.588–604.

Myers, C.D. et al., 1996. Developmental genetic analysis of troponin T mutations in striated and nonstriated muscle cells of Caenorhabditis elegans. The Journal of Cell Biology, 132(6), pp.1061–1077.

Nagai, T. & Brown, B.E., 1969. Insect visceral muscle. Electrical potentials and contraction in fibres of the cockroach proctodeum. Journal of Insect Physiology, 15(11), pp.2151–2167.

Nishida, W. et al., 2002. A Triad of Serum Response Factor and the GATA and NK Families Governs the Transcription of Smooth and Cardiac Muscle Genes. Journal of Biological Chemistry, 277(9), pp.7308–7317.

Nyitray, L. et al., 1994. Scallop striated and smooth muscle myosin heavy-chain isoforms are produced by alternative RNA splicing from a single gene. Proceedings of the National Academy of Sciences, 91(26), pp.12686–12690.

Nylund, A. et al., 1988. Heart ultrastructure in four species of Onychophora (Peripatopsidae and Peripatidae) and phylogenetic implications. Zool Beitr, 32, pp.17–30.

Obinata, T., Ooi, A. & Takano-Ohmuro, H., 1983. Myosin and actin from ascidian smooth muscle and their interaction. Comparative Biochemistry and Physiology Part B: Comparative Biochemistry, 76(3), pp.437–442.

Ojima, T. & Nishita, K., 1986. Isolation of Troponins from Striated and Smooth Adductor Muscles of Akazara Scallop. Journal of Biochemistry, 100(3), pp.821–824.

Økland, S., 1980. The heart ultrastructure of Lepidopleurus asellus (Spengler) and Tonicella marmorea (Fabricius)(Mollusca: Polyplacophora). Zoomorphology, 96(1–2), pp.1–19.

OOta, S. & Saitou, N., 1999. Phylogenetic relationship of muscle tissues deduced from superimposition of gene trees. Molecular Biology and Evolution, 16(6), pp.856–867.

Paniagua, R. et al., 1996. Ultrastructure of invertebrate muscle cell types. Histology and Histopathology, 11(1), pp. 181–201.

Phiel, C.J. et al., 2001. Differential Binding of an SRF/NK-2/MEF2 Transcription Factor Complex in Normal Versus Neoplastic Smooth Muscle Tissues. Journal of Biological Chemistry, 276(37), pp.34637–34650.

Philippe, H. et al., 2011. Acoelomorph flatworms are deuterostomes related to 24 Xenoturbella. Nature, 470(7333), pp.255–258.

Prosser, C.L., Nystrom, R.A. & Nagai, T., 1965. Electrical and mechanical activity in intestinal muscles of several invertebrate animals. Comparative biochemistry and physiology, 14(1), pp.53–70.

Raible, F. et al., 2005. Vertebrate-type intron-rich genes in the marine annelid Platynereis dumerilii. Science, 310(5752), pp.1325–1326.

Reynolds, P.D., Morse, M.P. & Norenburg, J., 1993. Ultrastructure of the Heart and Pericardium of an Aplacophoran Mollusc (Neomeniomorpha): Evidence for Ultrafiltration of Blood. Proceedings of the Royal Society of London B: Biological Sciences, 254(1340), pp.147–152.

Rieger, R. et al., 1991. Organization and differentiation of the body-wall musculature in Macrostomum (Turbellaria, Macrostomidae). Hydrobiologia, 227(1), pp.119–129.

Roach, D.K., 1968. Rhythmic muscular activity in the alimentary tract of Arion ater (L.)(Gastropoda: Pulmonata). Comparative biochemistry and physiology, 24(3), pp.865–878.

Rogers, D.C., 1969. Fine structure of smooth muscle and neuromuscular junctions in the foot of Helix aspersa. Zeitschrift für Zellforschung und Mikroskopische Anatomie, 99(3), pp.315–335.

Rosenbluth, J., 1972. Obliquely striated muscle. In The structure and function of muscle. New York and London: Academic Press, pp. 389–420.

Royuela, M. et al., 2000. Characterization of several invertebrate muscle cell types: a comparison with vertebrate muscles. Microscopy Research and Technique, 48(2), pp.107–115.

Royuela, M. et al., 1997. Immunocytochemical electron microscopic study and Western blot analysis of caldesmon and calponin in striated muscle of the fruit fly Drosophila melanogaster and in several muscle cell types of the earthworm Eisenia foetida. European journal of cell biology, 72(1), pp.90–94.

Sanders, K.M., Koh, S.D. & Ward, S.M., 2006. Interstitial cells of Cajal as pacemakers in the gastrointestinal tract. Annu. Rev. Physiol., 68, pp.307–343.

Saudemont, A. et al., 2008. Complementary striped expression patterns of NK homeobox genes during segment formation in the annelid Platynereis. Developmental biology, 317(2), pp.430–443.

Schlesinger, J. et al., 2011. The Cardiac Transcription Network Modulated by Gata4, Mef2a, Nkx2.5, Srf, Histone Modifications, and MicroRNAs. PLOS Genet, 7(2), p.e1001313.

Schmidt-Rhaesa, A., 2007. The Evolution of Organ Systems 1 edition., Oxford; New York: Oxford University Press.

Sebé-Pedrós, A. et al., 2014. Evolution and classification of myosins, a paneukaryotic whole-genome approach. Genome biology and evolution, 6(2), pp.290–305.

Shaw, K., 1974. The fine structure of muscle cells and their attachments in the tardigrade Macrobiotus hufelandi. Tissue and Cell, 6(3), pp.431–445.

Shi, X. & Garry, D.J., 2006. Muscle stem cells in development, regeneration, and disease. Genes & Development, 20(13), pp.1692–1708.

Shimeld, S.M. et al., 2010. Clustered Fox genes in lophotrochozoans and the evolution of the bilaterian Fox gene cluster. Developmental biology, 340(2), pp.234–248.

Silverthorn, D.U., 2015. Human Physiology: An Integrated Approach 7 edition., San Francisco: Pearson.

Spies, R.B., 1973. The blood system of the flabelligerid polychaete Flabelliderma commensalis (Moore). Journal of morphology, 139(4), pp.465–490.

Steinmetz, P.R.H. et al., 2012. Independent evolution of striated muscles in cnidarians and bilaterians. Nature, 487(7406), pp.231–234.

Sulbarán, G. et al., 2015. An invertebrate smooth muscle with striated muscle myosin filaments. Proceedings of the National Academy of Sciences, 112(42), pp.E5660–E5668.

Susic-Jung, L. et al., 2012. Multinucleated smooth muscles and mononucleated as well as multinucleated striated muscles develop during establishment of the male reproductive organs of Drosophila melanogaster. Developmental Biology, 370(1), pp.86–97.

Sweeney, H.L., Bowman, B.F. & Stull, J.T., 1993. Myosin light chain phosphorylation in vertebrate striated muscle: regulation and function. American Journal of Physiology-Cell Physiology, 264(5), pp.C1085–C1095.

Thor, S. & Thomas, J.B., 2002. Motor neuron specification in worms, flies and mice: conserved and “lost” mechanisms. Current Opinion in Genetics & Development, 12(5), pp.558–564.

Tjønneland, A., Økland, S. & Nylund, A., 1987. Evolutionary aspects of the arthropod heart. Zoologica Scripta, 16(2), pp.167–175.

Wagner, G.P., 2014. Homology, Genes, and Evolutionary Innovation, Princeton ; Oxford: Princeton University Press.

Wales, S. et al., 2014. Global MEF2 target gene analysis in cardiac and skeletal muscle reveals novel regulation of DUSP6 by p38MAPK-MEF2 signaling. Nucleic Acids Research, p. gku813.

Wang, D.-Z. & Olson, E.N., 2004. Control of smooth muscle development by the myocardin family of transcriptional coactivators. Current Opinion in Genetics & Development, 14(5), pp.558–566.

Wang, Z. et al., 2003. Myocardin is a master regulator of smooth muscle gene expression. Proceedings of the National Academy of Sciences, 100(12), pp.7129–7134.

White, J., 1988. 4 The Anatomy. The nematode Caenorhabditis elegans – Cold Spring Harbor Monograph Archive, 17, pp.81–122.

Witchley, J.N. et al., 2013. Muscle Cells Provide Instructions for Planarian Regeneration. Cell Reports, 4(4), pp.633–641.

Wood, J.D., 1969. Electrophysiological and pharmacological properties of the stomach of the squid Loligo pealii (Lesueur). Comparative biochemistry and physiology, 30(5), pp.813–824.

Wu, K.S., 1939. On the physiology and pharmacology of the earthworm gut. J. exp. Biol., 16, pp.184–197.

Zaffran, S. et al., 2001. biniou (FoxF), a central component in a regulatory network controlling visceral mesoderm development and midgut morphogenesis in Drosophila. Genes & Development, 15(21), pp.2900–2915.

Zhang, Y. et al., 2000. Drosophila D-titin is required for myoblast fusion and skeletal muscle striation. Journal of cell science, 113(17), pp.3103–3115.

Zhou, Q. et al., 2001. Ablation of Cypher, a PDZ-LIM domain Z-line protein, causes a severe form of congenital myopathy. The Journal of Cell Biology, 155(4), pp.605–612.

